# Surveying the landscape of RNA isoform diversity and expression across 9 GTEx tissues using long-read sequencing data

**DOI:** 10.1101/2024.02.13.579945

**Authors:** Madeline L. Page, Bernardo Aguzzoli Heberle, J. Anthony Brandon, Mark E. Wadsworth, Lacey A. Gordon, Kayla A. Nations, Mark T. W. Ebbert

**Author notes:** To whom correspondence should be addressed: Mark T.W. Ebbert.

## Abstract

Even though alternative RNA splicing was discovered nearly 50 years ago (1977), we still understand very little about most isoforms arising from a single gene, including in which tissues they are expressed and if their functions differ. Human gene annotations suggest remarkable transcriptional complexity, with approximately 252,798 distinct RNA isoform annotations from 62,710 gene bodies (Ensembl v109; 2023), emphasizing the need to understand their biological effects. For example, 256 gene bodies have ≥50 annotated isoforms and 30 have ≥100, where one protein-coding gene (*MAPK10*) even has 192 distinct RNA isoform annotations. Whether such isoform diversity results from biological redundancy or spurious alternative splicing (i.e., noise), or whether individual isoforms have specialized functions (even if subtle) remains a mystery for most genes. Recent studies by Aguzzoli-Heberle et al., Leung et al., and Glinos et al. demonstrated long-read RNAseq enables improved RNA isoform quantification for essentially any tissue, cell type, or biological condition (*e.g.,* disease, development, aging, etc.), making it possible to better assess individual isoform expression and function. While each study provided important discoveries related to RNA isoform diversity, deeper exploration is needed. We sought to quantify and characterize real isoform usage across tissues (compared to annotations). We used long-read RNAseq data from 58 GTEx samples across nine tissues (three brain, two heart, muscle, lung, liver, and cultured fibroblasts) generated by Glinos et al. and found considerable isoform diversity within and across tissues. Cerebellar hemisphere was the most transcriptionally complex tissue (22,522 distinct isoforms; 3,726 unique); liver was least diverse (12,435 distinct isoforms; 1,039 unique). We highlight gene clusters exhibiting high tissue-specific isoform diversity per tissue (*e.g., TPM1* expresses 19 in heart’s atrial appendage). We also validated 447 of the 700 new isoforms discovered by Aguzzoli-Heberle et al. and found that 88 were expressed in all nine tissues, while 58 were specific to a single tissue. This study represents a broad survey of the RNA isoform landscape, demonstrating isoform diversity across nine tissues and emphasizes the need to better understand how individual isoforms from a single gene body contribute to human health and disease.

**Dear reviewers:** We sincerely appreciate the time and effort you are taking to review our manuscript. We recognize it is a substantial commitment and welcome your feedback to ensure this work is accurate and helpful to furthering the field’s understanding of the human genome and its relevance to human health and disease. Because we recognize how important it is for all scientists to receive proper credit for their contributions to the field, **we specifically invite you to notify us if we failed to cite or give proper credit to any relevant publications, whether they be yours or another group’s work.** Of course, we also welcome all other feedback and will do our best to respond to your suggestions and concerns.

Sincerely,

Mark T. W. Ebbert

## Introduction

According to our analysis of Ensembl gene annotations v109 (released February 2023)^1^, 38% of human gene bodies (85% protein coding) express multiple RNA and protein isoforms via alternative splicing, corroborating other analyses^2,3^. Yet, despite researchers knowing about alternative splicing for decades^4–6^, little is known about individual isoforms for most genes, including if or how their functions differ. In fact, when discussing a given gene’s function, researchers often speak as if the gene has a single function, despite the many RNA and protein isoforms it expresses. We argue that one of the most important next-steps in understanding the biology of complex organisms, including human health and disease, will be to understand the functions of individual RNA and protein isoforms arising from a “single” gene (*i.e.,* expressed from a single gene body). While by definition, a gene is simply a specific DNA sequence, if a single gene body gives rise to multiple RNA and protein isoforms with different functions, we question whether it truly is a “single” gene in practice.

Historically, RNA sequencing studies have collapsed expression across all RNA isoforms for a given gene into a single gene expression measurement due to technical limitations in short-read sequencing, ignoring the underlying complexity. This approach prevents researchers from: (1) discovering and characterizing all isoforms for a given gene, (2) quantifying expression for distinct isoforms, and (3) fully understanding each isoform’s function, as it is not possible to truly understand an isoform’s function without knowing its expression patterns across all tissues and cell types.

Given the complexity of many eukaryotes, including humans, it is reasonable to assume that distinct RNA and protein isoforms from a single gene body may have unique functions, even if the functions are closely related. There are a few well-documented examples where different isoforms from a single gene body have clearly different (even opposite) functions. Perhaps the most recognized example is *BCL-X* (*BCL2L1*)^7^, where one isoform is pro-apoptotic (BCL-Xs) while the other is anti-apoptotic (BCL-XL). Additional examples, where different isoforms appear to have more subtle functional differences, include: (1) *RAP1GDS1* (also known as *SmgGDS*)^8^, where its isoforms interact differently with small GTPases^8,9^; and (2) *TRPM3*, which encodes cation-selective channels in humans, and can be alternatively spliced into two variants targeting different ions^10–12^. *RAP1GDS1* and *TRPM3* showcase isoforms performing functions that are closely related yet not identical, while *BCL-X* (*BCL2L1*) is an excellent example where the isoforms perform entirely opposite functions.

Given the advent of long-read sequencing technologies like PacBio and Oxford Nanopore Technologies (ONT), it is now possible to more accurately characterize and quantify expression for individual RNA isoforms expressed for each gene across essentially any tissue, cell type, or biological condition (*e.g.,* diseases, development, aging, etc.). Long reads are not perfect because of RNA degradation and technical challenges^13^, but along with work by Leung et al.^14^, our recent work in Aguzzoli-Heberle et al.^2^ demonstrated the value of long-read sequencing in human brain and the technology’s ability to quantify expression for individual RNA isoforms, including *de novo* isoforms and entirely new gene bodies.

Similarly, Glinos et al.^15^ performed long-read RNA sequencing across 15 tissues, providing valuable data to begin assessing expression patterns for individual RNA isoforms across human tissues. By characterizing and quantifying expression for individual RNA isoforms across human tissues, we can begin to understand whether different RNA isoforms from a single gene body perform different functions—even if subtle—where an isoform might be more carefully tuned for the tissue it is expressed in. In fact, some of the subtler differences may explain challenging medical mysteries that remain unresolved, including why patients react differently to certain exposures (*e.g.,* medicines, vaccines, etc.).

In their research, Glinos et al.^15^ initiated an investigation into tissue-specific RNA isoform expression, but given the broad scope of their article and the crucial importance of this subject, we believe a more thorough exploration is warranted. To provide a broad survey and to characterize the RNA isoform landscape across the human genome, and emphasize the importance of alternative splicing in complex organisms, including humans, our aims are three-fold; (1) briefly characterize the history of gene body and RNA isoform discovery since 2014 by comparing Ensembl^1^ gene annotations from 2014 to 2023 to assess changes over time; (2) characterize individual isoform expression across nine GTEx tissues (58 samples total) using data generated by Glinos et al.^15^; and (3) verify the existence and quantify the expression of new RNA isoforms, including those from new gene bodies, that we previously discovered in Aguzzoli-Heberle et al.^2222^ across the nine GTEx tissues. We further provide a user-friendly website to explore the RNA isoform expression data for the GTEx samples (https://ebbertlab.com/gtex_rna_isoform_seq.html) and for our own RNA isoform expression data from prefrontal cortex (https://ebbertlab.com/brain_rna_isoform_seq.html)^2^.

## Results

Here, we present results beginning with a broad overview of gene body and RNA isoform discovery from 2014 to 2023, followed by a more focused analysis of RNA isoform expression across nine tissue types, using long-read data generated by Glinos et al.^15^ combined with results from our recent study of deep long-read cDNA sequencing in human prefrontal cortex (Brodmann 9/46) in Aguzzoli-Heberle et al.^2^

### Methodological overview

For this study (see study design in **Figure 1**), we retrieved long-read cDNA data for GTEx samples from Glinos et al.^15^ sequenced using the Oxford Nanopore Technologies (ONT) minION. For consistency across data sets, we reanalyzed samples using our bioinformatics pipeline. Analysis included isoform quantification with Bambu^16^ using Ensembl^1^ v109 annotations with the addition of the 700 new isoforms discovered in Aguzzoli-Heberle et al.^2^. We only retained tissues with at least five samples from unique subjects in our analyses. We further removed technical replicates, samples with experimental conditions (*i.e.*, *PTBP1* knockdown), and samples with low total read counts (< 1,000,000 reads; **Figure 1**). One liver sample was removed because it clustered poorly during principal components analysis (PCA; **Supplemental Figure S1-2; Supplemental Table S1**). Ensembl annotations across a ten-year period (2014-2023, one per year) were downloaded for additional analyses, as discussed below (**Figure 1**; **Supplemental Table S2)**.

**Figure 1:**
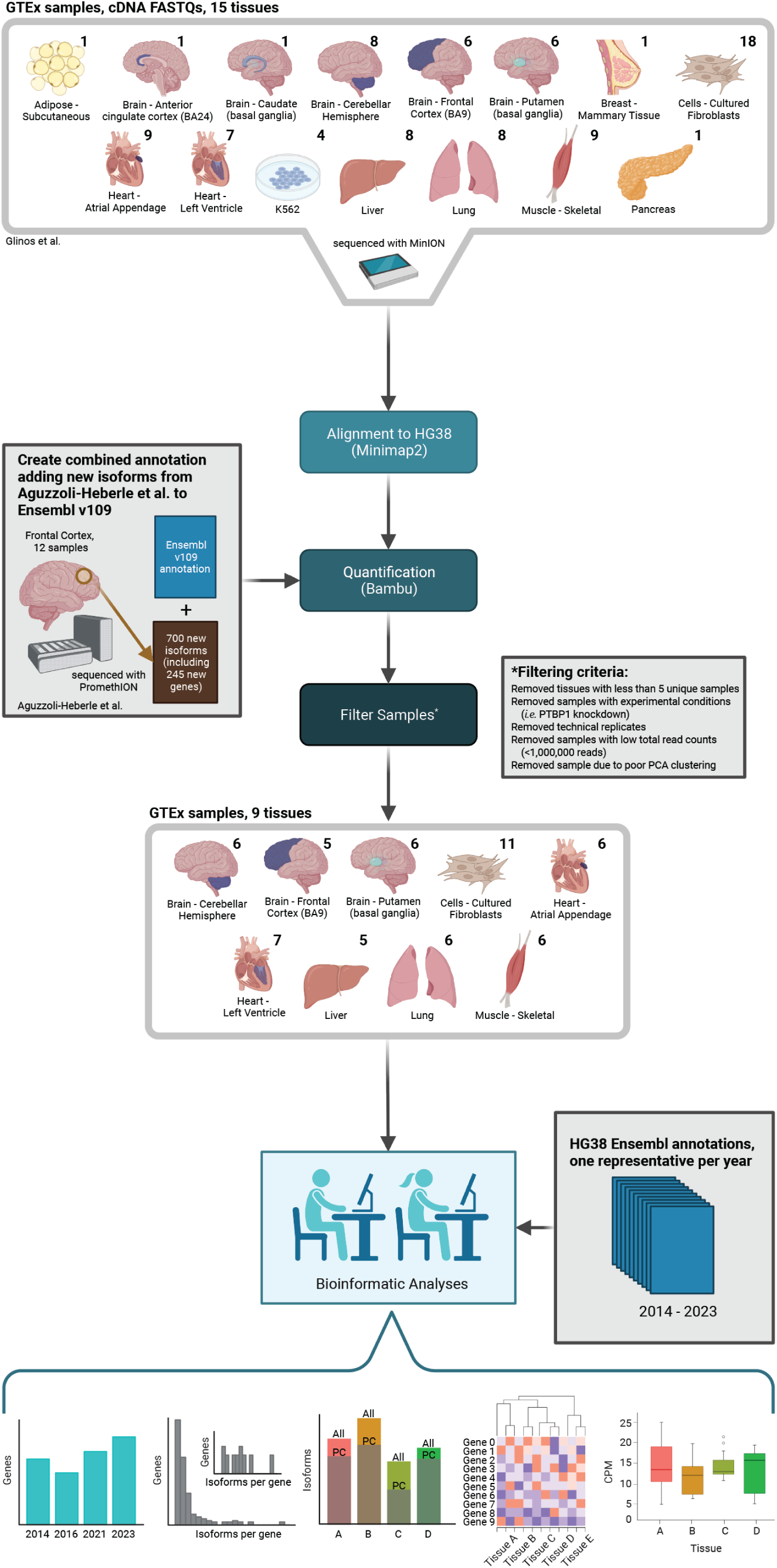
Study design to assess RNA isoform diversity across human tissues. We used long-read RNAseq data from GTEx samples generated by Glinos et al. to characterize and quantify RNA isoform expression and diversity across various tissues. These data were combined with results from our recent work in Aguzzoli-Heberle et al. GTEx data were reanalyzed for consistency across data sets, as shown. After filtering samples and tissues (as shown) we included nine tissues in our analyses. The numbers above each tissue indicate the number of samples. Created with BioRender.com.

While it is possible to sequence entire RNA isoforms with long-read sequencing, many reads still cannot be uniquely mapped to an isoform because of degradation and technical limitations^13^, even with high-quality RNA (*e.g.,* RIN ≥ 9). To address this challenge, Bambu—the quantification tool we employed—provides RNA isoform expression estimates in three forms, each with particular pros and cons: (1) total counts, (2) full-length counts, and (3) unique counts.

The primary advantage for the “total counts” metric is that it utilizes all aligned reads. Credit for a single read count is split for reads that map to multiple isoforms. For example, if a read is equally likely to originate from two isoforms, each isoform receives credit for 0.5 reads. The disadvantage is that this approach likely overestimates expression for some isoforms while underestimating expression for others, but is a valid approach given the challenge of assigning ambiguous reads to the correct isoform.

The “full-length counts” metric only includes reads containing all exon junctions for at least one isoform. If the read is still not unique to a single isoform, Bambu will split the read count among the relevant isoforms (same as total counts). Thus, the full-length counts metric reduces ambiguity, but also likely dramatically underrepresents longer isoforms because they will not be fully sequenced as often^13^. For reference, 51% of reads were “full-length” in this study.

The “unique counts” metric is the strictest and includes only reads that align *uniquely* to a single isoform, because of a unique exon-exon junction structure. In theory, unique counts would be the ideal metric because the alignment is unambiguous and would best represent the observed expression for all isoforms. Unfortunately, this approach is biased against isoforms that only differ towards the 5’ end (degradation bias) and isoforms that have the same exon-exon junctions but only differ in their start or end sites (e.g., shortened 3’ UTR).^16^. Approximately 63% of reads aligned uniquely to an isoform in this study; while far from perfect, this is still a significant improvement over short-read sequencing.

Currently, there is no perfect solution to the challenge of unambiguously assigning reads to the correct isoform. While we anticipate more sophisticated approaches will be developed in the coming years, we used a relatively simplistic approach involving both the total counts (to utilize all reads; *i.e.,* sensitivity) and unique counts (for specificity) provided by Bambu. Specifically, to minimize false positives, a given isoform was only included in this study if it had both a median counts-per-million (CPM) > 1 (total counts) and a median-unique-counts ≥ 1 across all samples within a given tissue, unless otherwise specified.

### Gene and RNA isoform annotation comparison from 2014-2023 reveals a general increase in gene body and RNA isoform discoveries and many annotated isoforms per gene

#### Gene body annotations

To quantify the rate of gene body and RNA isoform discoveries, we compared Ensembl annotations spanning 2014 to 2023 at both the gene and RNA isoform levels (**Figure 2**). Starting with Ensembl v76 (released August 2014), there were 58,764 gene annotations present, compared to 62,710 by Ensembl v109 (released February 2023; 3,946 increase; **Figure 2a**). We observed a sharp increase in new gene body annotations in 2015 (1,792), followed by a sharper decrease in 2016 (2,505). Between 2016 and 2023, there was a general increase in gene annotations, including large increases in 2020 and 2023. Comparing gene body annotations between 2019, 2021, and 2023, 196 and 14 were unique to 2019 and 2021, respectively, with 149 shared between them that were missing in 2023 (Venn diagram in **Figure 2a**), indicating that many gene body annotations were dropped between 2021 and 2023.

**Figure 2:**
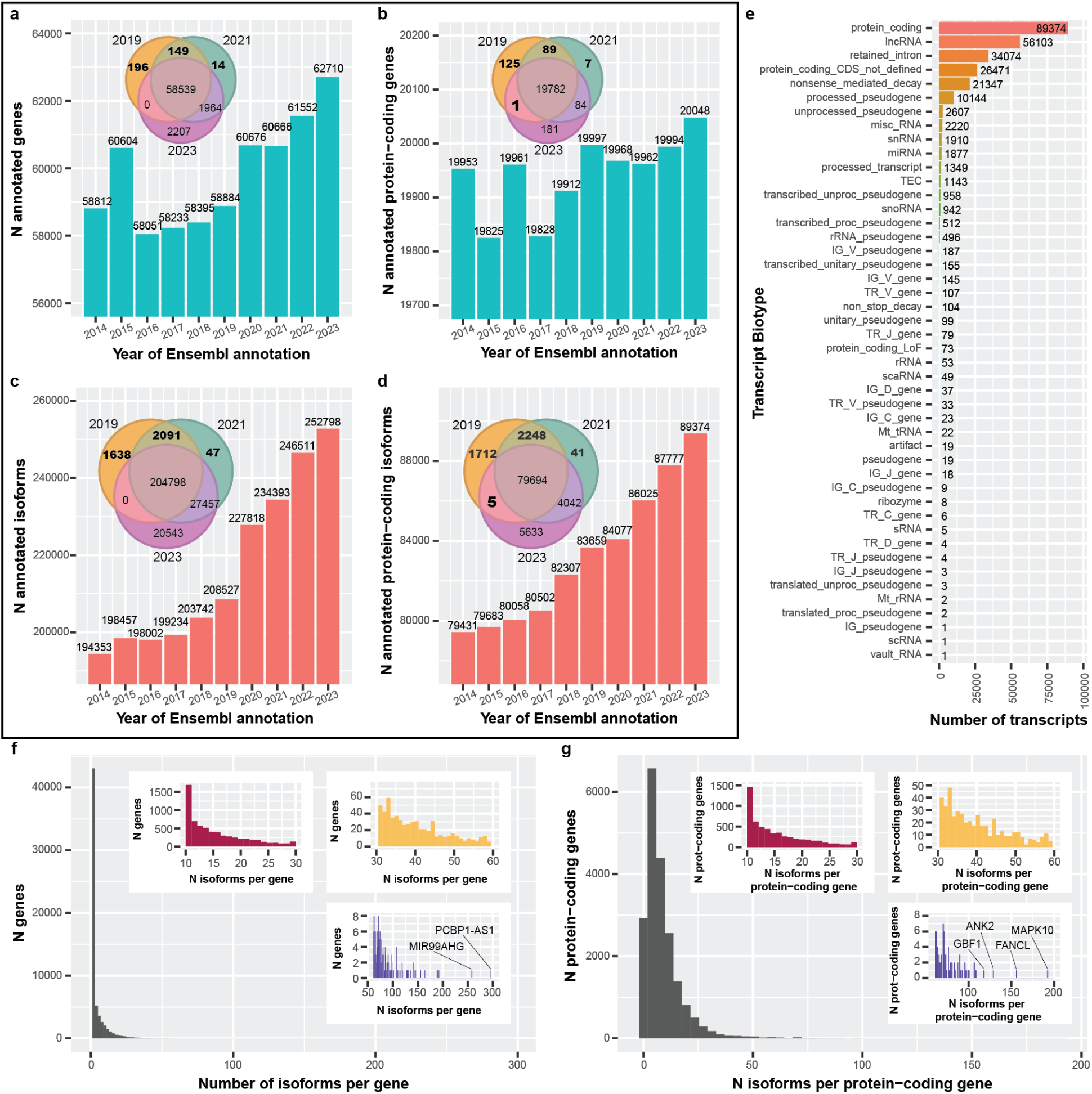
Ensembl gene and isoform annotations generally increased between 2014 & 2023, and reveal complex RNA isoform diversity. For figures (a-d), a representative Ensembl annotation was chosen for each year from 2014-2023. (a) The number of annotated genes per year from 2014-2023. After 2016, the overall trend is an increase, with a reasonably sharp increase around 2020. We see that some annotations appear to be dropped completely between years (Venn diagram). (b) Same as (a) but restricted to protein-coding genes. In contrast, there were more fluctuations in the number of annotated protein-coding genes, though still a positive trend. One gene was annotated as protein-coding in the 2019 and 2023 annotations but was not in 2021. (c) Looking specifically at RNA isoform annotations, the trend is positive, with a sharp increase in 2020. (d) Similar patterns exist for RNA isoforms annotated as protein-coding. There are five isoforms that were not labeled as protein-coding in 2021 but were in 2019 and 2023. (e) Bar plot showing transcript biotype for all isoforms in 2023. As expected, protein-coding was the most common biotype. Interestingly, retained intron was 3^rd^ and nonsense mediated decay was 5^th^. (f) Histogram showing number of annotated isoforms per gene. Colored, zoomed subplots shown for convenience. The majority of gene bodies had only one annotated isoform (median = 1; 75^th^ percentile: 4; 85^th^: 8; 95^th^: 16), but some had more than 100. The most annotated isoforms for a single gene body was 296 (PCBP1-AS1; ENSG00000179818). (g) Similar to (f) but showing the number of isoforms per protein-coding gene. The median number of annotated isoforms is 6 (75^th^ percentile: 11 isoforms). The most annotations for a single protein-coding gene body was 192 (MAPK10; ENSG00000109339).

Exactly why gene body annotations were dropped in subsequent releases is not clear, nor is it available in publicly accessible data (to our knowledge), but we point out the seeming inconsistencies to highlight how challenging annotating a genome is. Ensembl provides an ID History Converter tool that reports specific release versions whenever a gene ID had a major update. While useful for specific situations, the tool does not explicitly state what changed, nor could we access ID history via a high-throughput method (*i.e.*, it required manual curation). Our analysis does not account for gene ID changes because, to our knowledge, extracting this information through Ensembl is not currently possible via a high-throughput method. Documentation on why changes are made, and the ability to programmatically assess them genome-wide would provide increased transparency and improve scientific reproducibility. To be clear, however, Ensembl and their collaborators employ a manually supervised computational workflow combined with expert annotators to resolve these challenging problems^17^, and their contributions to the field are both significant and essential.

In contrast to the full set of gene body annotations, there were multiple large increases and decreases for protein-coding genes from 2014-2017 (**Figure 2b**) and small decreases between 2019 and 2021. Comparing protein-coding gene body annotations between 2019, 2021, and 2023, 125 and 7 were unique to 2019 and 2021, respectively, with 89 shared between them that were not present in 2023 (Venn diagram in **Figure 2b**). For interest, there was a single protein-coding gene body annotation present in 2019 and 2023 that was absent in 2021 (**Supplemental Table S3**); the annotation was present in all three releases, but the gene’s biotype changed from “protein coding” to “transcribed unprocessed pseudogene” in February 2021 (Ensembl release 103)^18^, but reverted to “protein coding” by December 2021 (Ensembl release 105)^19^ .

#### RNA isoform annotations

In our view, a reference genome is only as good as its annotations; thus, while having properly annotated gene bodies is critical, having fully characterized and annotated RNA isoforms for all genes is equally important for understanding an organism’s full complexity. Exactly 194,305 RNA isoform annotations existed in 2014 (Ensembl v76), compared to 252,798 in 2023 (58,493 increase); 19,291 were newly annotated between 2019 and 2020 alone (**Figure 2c**). We cannot verify why this sudden increase occurred, but we believe it is directly related to long-read sequencing technologies becoming more accessible in previous years. Comparing RNA isoform annotations between 2019, 2021, and 2023, 1,638 and 47 were unique to 2019 and 2021, respectively, with 2,091 shared between them that were not present in 2023, again showing annotations being dropped (Venn diagram in **Figure 2c**).

We saw similar patterns for RNA isoforms annotated as protein-coding, with 79,431 annotated in 2014 (Ensembl v76) and 89,374 in 2023 (Ensembl v109; 9,943 increase; **Figure 2d**). Comparing protein-coding RNA isoforms between 2019, 2021, and 2023, 1,712 and 41 were unique to 2019 and 2021, respectively with 2,248 shared between them that were not present in 2023 (Venn diagram in **Figure 2d**). Interestingly, there were 5 RNA isoforms annotated as protein-coding that were shared between 2019 and 2023 but were not annotated as protein-coding in 2021 (Venn diagram in **Figure 2d; Supplemental Table S3**).

We also characterized the transcript biotype for all RNA isoforms in Ensembl v109 (2023; **Figure 2e**). The top five annotation categories were protein-coding (with 89,374 RNA isoforms), lncRNA (56,103), retained intron (34,074), protein-coding CDS not defined (26,471), and nonsense mediated decay (21,347; **Figure 2e**).

#### RNA isoforms per gene

We quantified the distribution of RNA isoforms per gene body and, as expected, most gene bodies (38,690) have only one annotated isoform (median = 1; 75^th^ percentile: 4; 85^th^: 8; 95^th^: 16; **Figure 2f**). Comparatively, only 2,922 protein-coding genes (14.6%) have a single annotated isoform while the median number of annotated isoforms is 6 (75^th^ percentile: 11 isoforms), demonstrating the transcriptional complexity among protein-coding genes compared to non-coding genes. Remarkably, the most annotated isoforms for a single gene body and a single protein-coding gene were 296 (*PCBP1-AS1*; ENSG00000179818; **Figure 2f**) and 192 (*MAPK10*; ENSG00000109339; **Figure 2g**), respectively.

Similarly, we observed 7,255, 256, and 30 gene bodies with ≥10, ≥50, and ≥100 annotated isoforms, respectively, and 6,162, 169, and 94 protein-coding gene bodies for the same respective thresholds. While biology regularly surprises scientific expectations and humans are complex organisms, we were skeptical that any single gene body could legitimately express so many isoforms (*e.g.*, 50). Thus, we wanted to determine whether we actually observe genes expressing that many unique isoforms using the long-read sequencing data from the GTEx samples^15^ (described below).

### Long-read RNA isoform expression reveals high isoform diversity and varying protein-coding isoform ratios across nine tissues

#### RNA isoform diversity across tissues

With the large number of annotated RNA isoforms in Ensembl v109, we wanted to characterize and quantify individual isoforms being expressed in human tissues, relative to their annotations as described above. Using long-read RNAseq data from Glinos et al.^15^, we compared RNA isoform expression patterns across nine tissues, including three brain regions (cerebellar hemisphere, frontal cortex, and putamen), two heart regions (left ventricle and atrial appendage), liver, lung, skeletal muscle, and cultured fibroblast cells. We included cultured fibroblast cells for interest since many scientific experiments utilize them.

The number of distinct isoforms that exceeded our thresholds ranged between 12,435 (liver; from 8,857 genes) and 22,522 (cerebellar hemisphere; from 13,236 genes; **Figure 3a,b**)—a surprisingly large range (a difference of 10,087), demonstrating substantial molecular diversity across tissues. For interest, the total number of isoforms with a 0 < CPM ≤ 1 ranged from 5,691 (heart [left ventricle]) to 12,385 (cerebellar hemisphere; **Figure 3a**), demonstrating the large number of isoforms that were observed with a CPM ≤ 1. Using an ultra-conservative CPM threshold of 10, expressed isoforms ranged between 3,815 (liver) and 7,916 (cerebellar hemisphere; **Figure 3a**).

**Figure 3:**
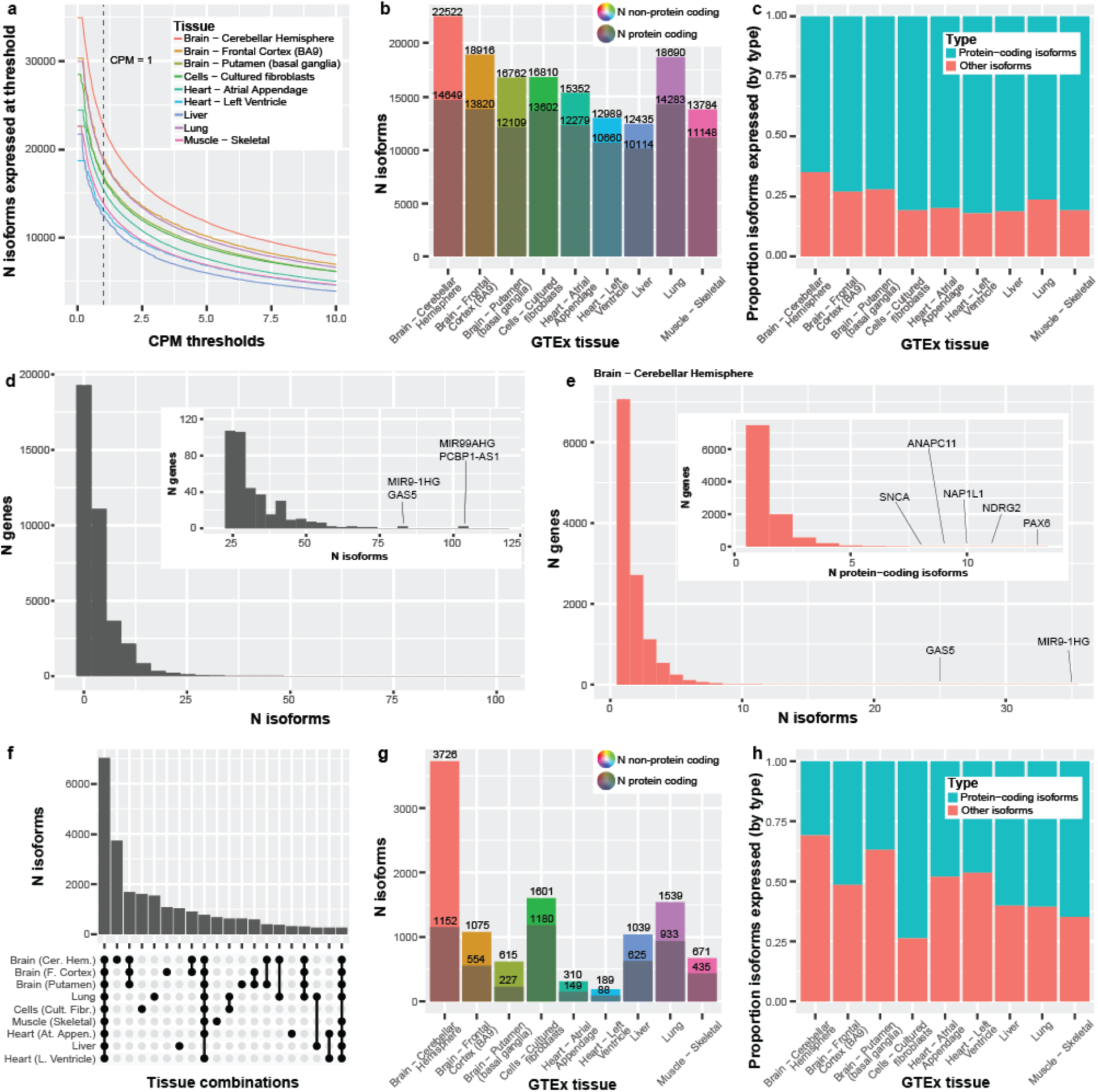
Long-read RNA isoform expression reveals high isoform diversity and variation across nine tissues. **(a)** Line graph showing the number of isoforms expressed across counts-per-million (CPM) thresholds from 0-10. Vertical line represents CPM = 1, where our threshold looks at everything with a CPM >1. The initial flat portion of each distribution is due to filtering out isoforms that did not have a median-unique-counts ≥ 1. **(b)** Number of isoforms and protein-coding isoforms (the darker shaded) expressed at our standard thresholds of median-unique-counts ≥ 1 and CPM > 1. The total number of expressed isoforms differs between tissues. **(c)** Bar plot comparing the proportion of protein-coding isoforms to other isoforms expressed across the nine GTEx tissues. Cerebellar hemisphere expressed the highest number of non-protein-coding RNA isoforms (7,873; 35.0%). Comparatively, the left ventricle of the heart expressed the smallest ratio of non-protein-coding isoforms (17.9%; 2,329 isoforms; Chi-square p = 1.90e-255). **(d)** The number of distinct isoforms expressed per gene, using a minimum threshold of one unique count in any sample for any tissue (maximum sensitivity). Zoomed subplot shown for conviencence. *PCBP1-AS1* expressed the most isoforms (105) for a single gene body and *FANCL* expressed the most (60) for a single protein-coding gene. 88.1% of gene bodies expressing ≥ 10 distinct RNA isoforms were protein coding, and 1,085 were medically relevant genes, as defined by Wagner et al.^20^, including PSEN2 (19 isoforms), PAX6 (43), and MAPK10 (46). Only ten gene bodies expressed ≥ 60 distinct RNA isoforms. **(e)** The number of isoforms per gene or number of protein-coding isoforms per gene (nested plot) expressed in cerebellar hemisphere using our standard threshold (median unique count ≥ 1 & CPM > 1). We saw 25 genes expressing ≥ 10 isoforms and 123 genes expressing ≥ 5 protein-coding isoforms. **(f)** Upset plot showing the first 20 interactions of isoform overlap between tissues. Exactly 7,023 isoforms had shared expression across all nine tissues. Cerebellar hemisphere has the largest number of isoforms uniquely expressed in a single tissue—more than half (53.1%) of the number of isoforms expressed across all nine tissues. **(g)** Bar plot showing the total number of isoforms that were uniquely expressed in each tissue. The grayed portion is the number of unique isoforms labeled protein-coding. **(h)** Bar plot comparing the proportion of protein-coding to non-protein-coding RNA isoforms, uniquely expressed by tissue. The cerebellar hemisphere expressed 3,726 isoforms uniquely, where only 30.9% (1,152) were protein-coding. Exactly 64.8% (435 out of 671) were protein-coding for skeletal muscle, however.

For RNA isoforms annotated as protein coding (regardless of the gene annotation), the number ranged from 10,114 in liver to 14,649 in cerebellar hemisphere (**Supplemental Figure S3**, **Figure 3b**). Isoforms expressed with a 0 < CPM ≤ 1 ranged from 2,888 (heart [left ventricle]) to 4,723 (cerebellar hemisphere; **Supplemental Figure S3**). Between 3,521 (liver) and 6,560 (cerebellar hemisphere) protein coding isoforms were expressed using an ultra-conservative threshold of 10 (**Supplemental Figure S3**). As expected, the top five (5) expressed transcript biotypes for each tissue closely mirrored the distributions from Ensembl annotations, with some minor exceptions (compare **Figure 2e** and **Supplemental Figure S4**).

Comparing the number of RNA isoforms expressed across tissues, the cerebellar hemisphere not only expressed substantially more total isoforms, but also a substantially larger proportion of non-protein-coding RNAs when compared to heart (**Figure 3b,c**). Specifically, cerebellar hemisphere expressed 7,873 (35.0%) non-protein-coding RNA isoforms compared to 2,329 for the heart’s left ventricle (17.9%; p = 1.90e-255; Chi-square test; **Figure 3b,c**); this includes all RNA isoforms not annotated as “protein coding”, even if it derived from a known protein-coding gene body. It is unclear whether the significant increase in non-protein-coding RNAs within cerebellar hemisphere compared to an arguably less complex tissue like the heart is biologically meaningful, but it merits further investigation; determining what role non-coding RNAs play in human health and disease remains a critical question in human biology.

#### RNA isoforms per gene

As discussed, Ensembl v109 suggests that 7,255, 256, and 30 genes express ≥ 10, ≥ 50, and ≥ 100 RNA isoforms, where the highest number of annotated isoforms for a single gene body was for *PCBP1-AS1* (lncRNA; 296) and, for protein-coding, *MAPK10* (192). We were skeptical that such a high number of isoforms could legitimately exist for a single gene. Thus, we quantified the number of isoforms we observed for all genes across the nine tissues included in this study.

To estimate the maximum number of distinct RNA isoforms we observed for each gene body across the nine tissues (erring on the side of sensitivity), we used a minimum threshold of one unique count in any sample for any tissue (*i.e.,* a single read that uniquely aligns to the given isoform from ≥ 1 sample). A large number of annotated isoforms for many of these genes are, in fact, observed at this forgiving threshold (**Figure 3d**; **Supplemental Table S4**). The most RNA isoforms observed for a single gene body and protein-coding body were 105 (*PCBP1-AS1*) and 60 (*FANCL*), respectively. Surprisingly, 2,751 of the 3,121 (88.1%) gene bodies expressing ≥ 10 distinct RNA isoforms were protein coding, and 1,085 were medically relevant genes, as defined by Wagner et al.^20^, including *PSEN2* (19 isoforms), *PAX6* (43), and *MAPK10* (46; **Supplemental Table S4-5**). Similarly, 231 of the 304 (80.0%) gene bodies expressing ≥ 25 distinct RNA isoforms were protein coding. We only observed ten gene bodies expressing ≥ 60 distinct RNA isoforms, where only one (10%) was protein coding. Considering that RNA expression measurements are simply a snapshot of expression in time, and the large number of additional tissues not included in this data set, it seems reasonable that some genes may legitimately express the additional RNA isoforms not observed here.

When using a stricter threshold of five unique counts, we observed 1,188 total gene bodes that expressed ≥ 10 distinct RNA isoforms, where 1,036 (87.2%) were protein coding (**Supplemental Table S5; Supplemental Figure S5a**); only three gene bodies expressed ≥ 60 RNA isoforms (all lncRNAs). Similarly, when using thresholds of 10 and 20 unique counts (**Supplemental Table S5; Supplemental Figure S5b,c**), we observed 608 (524 protein coding; 86.2%) and 270 (227 protein coding; 84.1%) gene bodies expressing ≥ 10 RNA isoforms, respectively; exactly two and one gene bodies expressed ≥ 60 RNA isoforms, respectively.

Returning to our standard and stricter threshold (median unique count ≥ 1 and CPM > 1), we quantified how many distinct isoforms per gene for a given tissue are consistently expressed. We still consistently observed gene bodies expressing ≥ 10 isoforms and protein-coding gene bodies expressing ≥ 5 isoforms at these high levels (**Figure 3e**; **Supplemental Figure S6**; **Supplemental Table S6**). Using cerebellar hemisphere as an example (**Figure 3e**), 25 genes expressed ≥ 10 isoforms and 123 expressed ≥ 5 protein-coding isoforms. Cerebellar hemisphere expresses more isoforms than others.

#### RNA isoforms per tissue and overlap

Looking at the overall number of expressed isoforms per tissue is itself informative, but understanding the overlap of isoforms expressed across tissues is equally important. Knowing which isoforms are common across all tissues versus those expressed in only a few or even a single tissue provides additional understanding about the potential function of the isoforms. Using our thresholds, 7,023 isoforms had shared expression across all nine tissues (**Figure 3f**). Cerebellar hemisphere uniquely expressed 3,726 isoforms—more than half (53.1%) of the number of isoforms expressed across all nine tissues, 2.22x more than the next largest overlap (1,680 isoforms; expressed in all three brain regions), and 19.7x more than the tissue with the least number of uniquely expressed isoforms (189; heart [left ventricle]; **Figure 3f,g**). These stark differences mark the cerebellar hemisphere as a truly unique tissue in this study, as similarly suggested by Glinos et al.^15^.

Curiously, the proportion of protein-coding isoforms expressed uniquely within a tissue varied. The cerebellar hemisphere uniquely expressed 3,726 isoforms, where only 30.9% (1,152 of 3,726) were protein-coding. Approximately 64.8% (435 out of 671) were protein-coding for skeletal muscle, however (**Figure 3g,h**). Cultured fibroblasts had the highest percentage of protein-coding isoforms at 73.7% (1,180 out of 1,601), though whether this would generalize to fibroblasts *in vivo*, we cannot say. Recent work by Cadiz et al.^21^, however, demonstrated that microglial cell cultures exhibited significantly unique expression signatures compared to freshly isolated microglia, demonstrating that cell cultures likely do not accurately represent reality. The large range in ratio of protein-coding isoforms to all other RNA isoforms solidifies our stance that understanding the interaction of isoforms in different tissues is needed.

#### Genes selectively express many isoforms per tissue

We were fascinated to find certain genes actively transcribe so many distinct isoforms in a single tissue type, which raises the question of why a single gene body is expressing multiple distinct isoforms simultaneously. Ultimately, targeted experimental work will be required to determine what function, if any, each isoform from a single gene body performs, but characterizing which genes and tissues these events occur in is the first step to understanding this phenomenon.

We selected all genes expressing > 5 distinct isoforms above our thresholds in at least one tissue, totaling 416 genes. We generated a clustered heatmap based on the isoform diversity (*i.e.,* the number of isoforms, not expression values) to identify gene clusters that preferentially express many isoforms for specific tissues (**Figure 4a**; **Supplemental Figure S7; Supplemental Table S7**). We identified several tissue-specific clusters, including for brain (as a whole), cerebellar hemisphere, muscle (including heart and skeletal muscle), liver, and lung (**Figure 4a**; **Supplemental Figure S7; Supplemental Table S7**). There are several fascinating examples of isoform complexity for each cluster, but we will only highlight a few.

**Figure 4:**
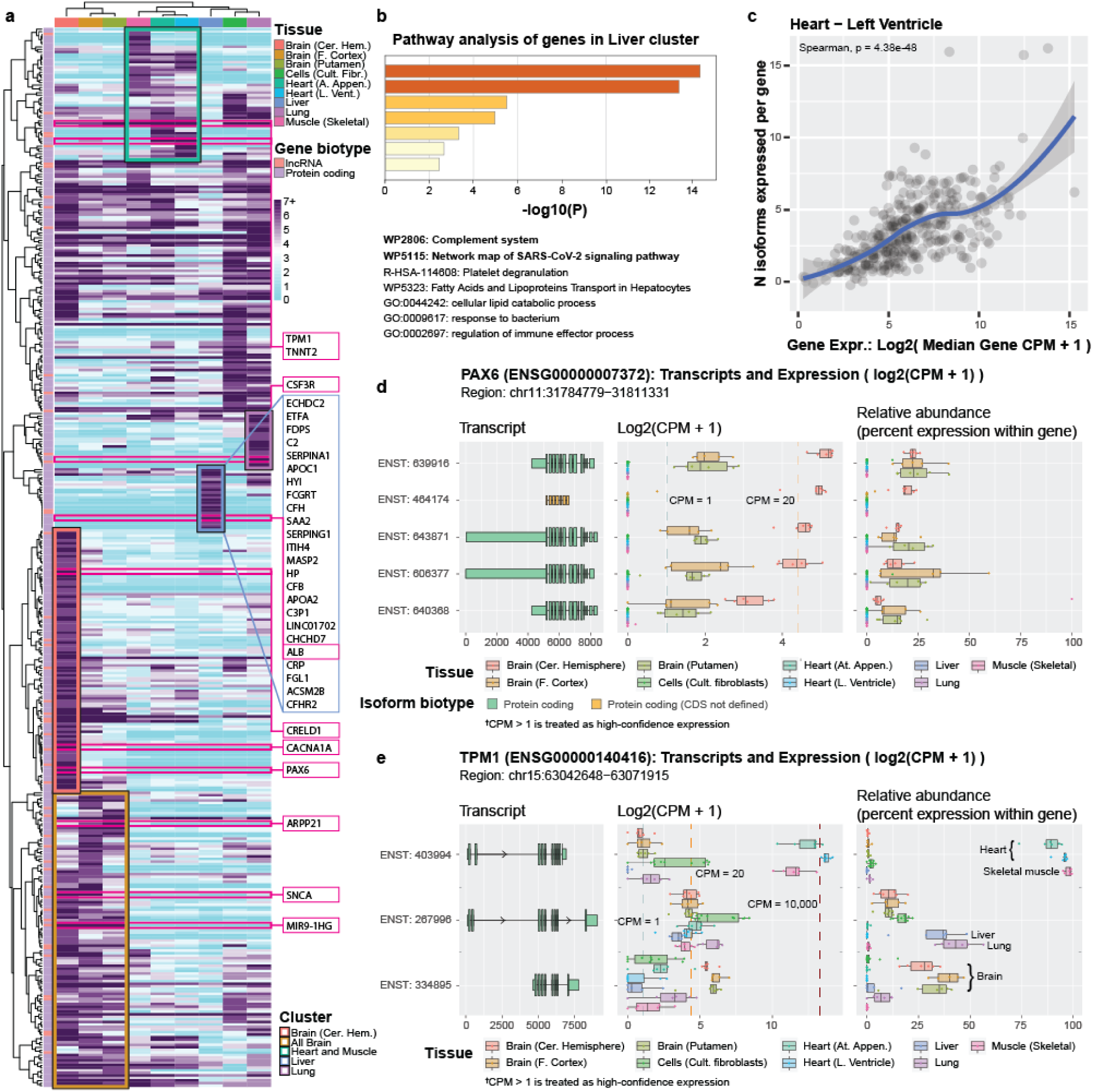
Genes selectively express many isoforms per tissue. We selected all genes expressing >5 isoforms in ≥1 tissue**. (a)** Clustered heatmap showing isoform diversity (i.e., number of isoforms, not expression values) to identify gene clusters that preferentially express many isoforms in specific tissues. Dark purple indicates ≥7 distinct isoforms. Each tissue set had a representative cluster. One gene (*MIR9-1HG*) expressed 36 isoforms in putamen. **(b)** Metascape pathway analysis for liver cluster shows enrichment for genes associated with immune response, including the complement system and SARS-CoV-2 signaling pathway. **(c)** Total gene expression and the number of isoforms expressed were correlated; left ventricle shown as representative sample (Spearman’s p = 4.38e-48; see **Supplemental Figure 18** for other tissues). We cannot determine cause and effect from these data alone, however, but can only conclude the two are correlated. Figures **(d-e)** show two representative gene examples from our R Shiny app (https://ebbertlab.com/gtex_rna_isoform_seq.html) expressing many isoforms within a single tissue, but with distinct expression patterns. For both genes (*PAX6* & *TPM1*), it is unclear whether overall gene expression is driving the high number of expressed isoforms or if the need for multiple isoforms causes a higher gene expression, but demonstrates the need to determine whether individual isoforms are biologically functional. **(d)** *PAX6* expressed 16 isoforms in cerebellar hemisphere; we highlight the top 5, here (see full plot in **Supplemental Figure 8**). **(e)** *TPM1* expressed 16 and 19 isoforms in left ventricle and atrial appendage, respectively. We highlight three *TPM1* isoforms showing preferential isoform expression across distinct tissue sets (see full plot in **Supplemental Figure 14**).

#### ARPP21, SNCA, and MIR9-1HG exhibit high isoform diversity in brain

An interesting example of RNA isoform diversity across all three brain regions, compared to the other tissues, is *ARPP21*, where little is known about this gene. *ARPP21* is highly expressed across brain regions and has been associated with entorhinal cortex thickness in an Alzheimer’s disease study^22^. It also displays high isoform diversity across all three brain regions included in this study, ranging from four (cerebellar hemisphere) to nine (putamen; **Figure 4a**; **Supplemental Figure S11; Supplemental Table S7**). Interestingly, the only other tissue within this study where any *ARPP21* isoform exceeds our threshold is skeletal muscle, which raises another critical point about gene and isoform function: given the distinct differences between the two tissues, what common molecular function fulfilled by *ARPP21* is needed by both brain and skeletal muscle but not other tissues (**Figure 4a**; **Supplemental Figure S8; Supplemental Table S7**). Indeed, according to GTEx, relative expression of *ARPP21* is highest in brain, followed by skeletal muscle, where most other tissues have very low expression^23^.

*SNCA*, acknowledged for its role in both Parkinson’s^24^ and Alzheimer’s disease^24–27^, is also highly expressed across all included brain regions and has elevated isoform diversity (putamen: 4; frontal cortex: 8; cerebellar hemisphere: 8) compared to other tissues that express this gene (lung: 2; heart [left ventricle]: 1; heart [atrial appendage]: 1; **Figure 4a**; **Supplemental Figure S9; Supplemental Table S7**). The transcript ENST00000394991 is expressed across all tissues that express *SNCA* and accounts for more than half of the total gene expression, but its expression is at least 5.5x higher in the three brain regions compared to other tissues, with median CPMs > 50 for all three regions. This isoform shares identical protein-coding sequence with four other isoforms expressed in this data (differing only in their untranslated regions [UTRs]), all of which have much lower CPM values (1.42 to 38.87 in brain). Why ENST00000394991 is clearly favored over the other isoforms when the only differences are in the UTR, and whether the UTR differences are biologically meaningful remain important biological questions.

As a non-protein-coding example, we observed a long non-coding RNA (lncRNA), *MIR9-1HG* (also known as *C1orf61*) that is only expressed in brain regions and is expressing between 32 and 36 distinct isoforms (**Figure 4a**; **Supplemental Figure S10**). While little is generally known about non-protein-coding gene bodies, they constitute 68% of all annotated gene bodies in Ensembl v109^1^. *MIR9-1HG*, specifically, has been implicated in development of ganglionic eminences, according to Zhao et al.^28^, which is consistent with *MIR9-1HG* being so highly expressed in brain tissue. Why *MIR9-1HG* expresses >30 distinct isoforms and whether they have distinct functions (or any function at all) is an important question.

#### PAX6, CACNA1A, and CRELD1 exhibit high isoform diversity in cerebellar hemisphere

We have highlighted that the cerebellar hemisphere has greater isoform diversity than other tissues, including other brain regions. Thus, it is not surprising that cerebellar hemisphere has its own cluster that is distinct from the other brain regions. (**Figure 4a; Supplemental Figure S11**). Two examples of isoform diversity within the cerebellar hemisphere are *PAX6* and *CACNA1A*, where only the three brain regions have any isoforms above our noise threshold. Only two to three isoforms met our criteria for putamen and prefrontal cortex for both genes, yet 16 and 11 isoforms met our criteria for the cerebellar hemisphere for *PAX6* and *CACNA1A*, respectively (**Figure 4a**; **Supplemental Figure S7,11,12; Supplemental Table S7**). Notably, *PAX6* is historically associated with eye diseases^29–31^ and early neural development^32^, yet it is already known to be highly expressed in the cerebellar hemisphere in adults^33^, suggesting a major function in this tissue. *CACNA1A*, on the other hand, is known to be directly involved in spinocerebellar ataxia type 6 (SCA6)^34^, where several DNA variants directly cause the disease. Thus, knowing the individual functions of individual RNA isoforms could be essential to treating this disease.

Similarly, the number of unique *CRELD1* isoforms ranges from three to five for all tissues except for the cerebellar hemisphere, where 12 exceeded our noise threshold (**Figure 4a**; **Supplemental Figure S13; Supplemental Table S7**)— double that of the next highest (liver). *CRELD1* is a known heart disease gene^35,36^, yet it has recently been implicated in a range of neurodevelopmental disorders, including epilepsy and movement disorders, for which the cerebellar hemisphere is major player^37^. A foundation dedicated to *CRELD1*-related neurodevelopmental diseases and associated research (CRELD1 Warriors; https://www.creld1.com/) has even been established in recent years^38^. Notably, total *CRELD1* expression is approximately three times higher in cerebellar hemisphere than in heart, according to GTEx^39^, and its isoform diversity in cerebellar hemisphere is four times greater (twelve) than what we observed in heart (three). The stark difference in isoform diversity between tissue regions of the same primary organ (*e.g.*, brain) is an excellent example demonstrating the need to understand why a single gene body can express many unique isoforms.

#### TPM1 and TNNT2 exhibit high isoform diversity in heart and skeletal muscle

According to the GTEx portal^40^, *TPM1* has high expression across several tissues, including heart and skeletal muscle, and plays an important role in cellular structure across numerous cell and tissue types. In muscle cells, specifically, *TPM1* provides structural support to the actin filament^41^. Most of the nine tissues expressed between two and nine isoforms (**Figure 4a**; **Supplemental Table S7**), but 16 and 19 RNA isoforms were expressed in the heart’s left ventricle and atrial appendage, respectively, demonstrating *TPM1*’s heart-specific isoform diversity. Total gene expression was a median CPM of 14,355.69 and 8,890.88, respectively, including isoforms that did not exceed our noise threshold (**Figure 4a**; **Supplemental Figure S14**; **Supplemental Table S7**). Those expression values are roughly 1,344x and 859x greater than the total median expression value in each tissue, respectively. The tissue expressing the next highest number of distinct isoforms was cultured fibroblasts with eleven, but whether this pattern would generalize to fibroblasts *in vivo* is unknown.

The *TNNT2* gene is directly associated with various cardiomyopathies^42–44^ and is primarily expressed in heart muscle^45^. Like *TPM1*, *TNNT2* expresses a large repertoire of RNA isoforms (16), but six are annotated as non-protein-coding (**Figure 4a**; **Supplemental Figure S15**; **Supplemental Table S7**)—again raising the question regarding what role each of the individual isoforms play in heart health. The complexities of understanding how genes interact throughout tissues are staggering. Adding the complexity of isoform diversity is even more difficult to comprehend. Interestingly, five *TNNT2* isoforms were expressed in prefrontal cortex and one in lung , but those are expressed at much lower levels (median CPMs ranged from 1.39 to 29.5) than what we observed for TNNT2 in heart (median CPMs ranged from 1.01 to 3,494.74 for the 16 isoforms; **Figure 4a**; **Supplemental Table S7**). It is interesting that the most highly expressed isoform in frontal cortex is not the isoform that is most highly expressed in heart. Whether the relatively lowly expressed isoforms in prefrontal cortex and lung are performing important biological functions in these tissues remains unknown.

#### Albumin (ALB) isoform diversity in liver may explain its many known functions

The albumin protein (ALB) was allegedly one of the first proteins discovered, and one of the most studied in history, where it was first precipitated from urine circa 1500 A.D.^46^. The breadth of ALB’s functions is remarkable. The ALB protein is expressed and excreted by the liver and is widely reported to be the most abundant protein in blood plasma^46–49^. ALB is a multifunctional protein that contributes approximately 70% of osmotic pressure^47,50^ while also serving to bind and transport a range of endogenous and exogenous compounds, including reactive oxygen species (ROS), pharmaceuticals, and hormones^46–50^. We observed eight *ALB* isoforms expressed in the liver (**Figure 4a; Supplemental Figure S16; Supplemental Table S7**), seven of which are annotated as protein-coding with distinct CDS regions. No other tissue expresses a single *ALB* isoform passing our thresholds. The most highly expressed isoform is ENST00000295897 (median CPM: 31,327.14), followed by ENST00000415165 (median CPM: 7,896.24). For reference, the median CPM for all isoforms expressed in the liver is 4.51, even when excluding isoforms that did not reach our noise threshold. Given the many functions attributed to *ALB*, it seems plausible that the multiple isoforms afforded by alternative splicing make this possible. Interrogating these isoforms further would allow us to determine what the differences are in the protein products and if the proteins have different or complementary functions.

Using the 24 genes in the liver cluster (**Figure 4a**), we performed a pathway enrichment analysis using Metascape^51^ and found enrichment for immune response, including the “complement system” and “network map of SARS-CoV-2 signaling pathways” (**Figure 4b**).

#### CSF3R isoform diversity in lung highlights lung’s role in immunity

A part of the lung cluster, *CSF3R* is a gene that encodes a cytokine receptor and is involved in the creation and regulation of granulocytes, a type of white blood cell. *CSF3R* has been directly implicated as a causal gene for chronic neutrophilic leukemia and atypical chronic myloid leukemia^52,53^, and has also been associated with neutropenia (insufficient neutrophils)^54^. In all, *CSF3R* is strongly implicated as an immune-related gene, and neutrophils specifically, yet we observed seven different isoforms above our thresholds in the lung, while other tissues expressed zero or one isoforms (**Figure 4a; Supplemental Figure S17**). Observing a neutrophil-related gene expressing seven distinct isoforms in the lungs may seem counterintuitive, initially, but the lungs host a large number of both innate and adaptive immune cells to protect against the regular exposure to pathogens^55,56^. Of the seven observed isoforms, three are annotated as protein-coding, three are annotated as retained intron, and one is protein-coding without a defined coding sequence (CDS). For *CSF3R*, the differences between protein-coding isoforms appear to be minimal compared to some of the other genes we have highlighted, raising the question of what the biological importance of each isoform is. Larger studies, deeper analyses, and ultimately experimental work will be required to determine the functional changes and their importance.

#### A word of caution: total gene expression and number of expressed isoforms are correlated

While we have highlighted gene clusters that express an enriched number of isoforms in specific tissues, it is important to exercise caution when drawing conclusions. In biology, proper interpretation is often not simple. There are likely additional underlying reasons for these genes to express multiple isoforms in a single tissue and not in others. For example, it is possible that the increased number of isoforms exceeding our threshold results from overall increased gene expression and potentially spurious alternative splicing. It is also important not to conflate isoform diversity and complexity with total expression. Even though a given gene body may be expressing ten discrete isoforms, most of the expression generally comes from one or two specific isoforms. Still, some of the isoforms with lower expression relative to other isoforms from the same gene body are expressed at levels much higher than most genes as a whole. Understanding the biological purpose in isoform diversity is, indeed, complex.

To assess whether increased isoform counts could result from increased gene expression, we tested for a correlation between the two measures using Spearman correlation and found that total gene expression is, in fact, correlated with the number of distinct isoforms expressed (p = 4.38e-48 for heart [left ventricle]; **Figure 4c**, **Supplemental Figure S18**), appearing to support the hypothesis that the number of observed isoforms may be driven by overall gene expression— at least in part. On the other hand, overall gene expression would also increase if a given cell is intentionally expressing multiple distinct isoforms. Thus, we cannot infer cause or biological significance from these data alone, but can only conclude that the two metrics are correlated. We think it is likely that some isoform diversity is caused by spurious splicing events, but that much is likely functional. Specifically, it has long been established that alternative splicing is an evolved process that enables biological diversity and complexity^57–59^ and it seems unlikely to us that only 20,000+ protein-coding genes performing a single function could support organisms as complex as humans with such diverse cell types, tissues, and developmental stages. Regardless of whether the distinct isoforms expressed from a single gene perform distinct functions (even if subtle), we need to fully characterize their nature to truly understand the underlying biology of human health and disease.

Here, we provide two examples (*PAX6* & *TPM1*) with different expression patterns, where it is unclear if total gene expression is driving the number of isoforms or if the number of isoforms is driving total gene expression. As discussed, *PAX6* expresses 16 isoforms above our thresholds in cerebellar hemisphere (**Supplemental Figure S8**). Twelve of the 16 isoforms represent only minimal relative expression, ranging from 0.71% to 4.96%, the biological significance of which is unclear. The other four, however, have high expression (CPM between 19.99 and 35.67) and arguably similar relative expression levels (ranging from 12.69% to 22.22%; **Figure 4d**). Given the top four isoforms constitute 68.74% of total *PAX6* expression with similar relative abundances, it could indicate the four isoforms are specifically necessary for optimal cellular function. In other words, based on these data alone, we cannot draw clear conclusions about the biological necessity of the twelve lowest-expressed isoforms within cerebellar hemisphere, nor can we determine the biological function of the top four isoforms. We can say, however, that at least four isoforms are actively expressed, and thus deeper work is needed to formally assess their function.

*TPM1* also appears to raise the same question (is overall gene expression driving isoforms expression or vice versa) but based on a different expression pattern compared to *PAX6*. As discussed, total *TPM1* expression levels in the heart’s left ventricle and atrial appendage are extremely high, being 1,344x and 859x greater than the median expression value in the respective tissues (10.68 and 10.35). Thus, given such high expression, it is plausible that many of the 16 and 19 expressed isoforms may have met our inclusion criteria due to potential spurious splicing. Additionally, a single isoform (ENST00000403994) constitutes between 89.2% and 98.2% of *TPM1*’s total expression for skeletal muscle and the two heart tissues (next highest tissue at 2.0%; **Figure 4e**; **Supplemental Figure S14**).

The evidence supporting at least some of the additional *TPM1* isoforms as biologically functional, however, is that three isoforms appear to have tissue-specific expression, where ENST00000403994, ENST00000267996, and ENST00000334895 are preferentially expressed in heart and skeletal muscle, lung and liver (percent abundance: 37.1% to 43.0%; next highest tissue at 18.3% ; **Figure 4e**), and brain tissues (percent abundance: 28.1% to 40.1%; next highest tissue at 6.8%; **Figure 4e**), respectively. Whether these distinct isoforms are ultimately functionally different remains to be seen. In either case, the examples of *PAX6* and *TPM1* highlight the need to exercise caution when interpreting results, as individual metrics can be misleading.

### New between 2019 and 2023

#### New RNA isoforms between 2019 and 2023, including isoforms from medically relevant genes

Many RNA isoforms were discovered between April 2019 (Ensembl v96) and February 2023 (Ensembl v109; 44,271), where 19,291 were newly annotated between 2019 and 2020, alone (**Figure 2c**). To assess the value added in those four years of RNA isoform discovery, we quantified the expression of new isoforms within the nine GTEx tissues. Here, we excluded isoforms recently discovered in Glinos et al.^15^, Leung et al.^14^, and Aguzzoli-Heberle et al.^2^. Using our threshold, the number of RNA isoforms discovered between 2019 and 2023 expressed in individual tissues ranged from 594 (liver) to 2,054 (cerebellar hemisphere; **Figure 5a**), including 180 and 508 from medically relevant genes (**Figure 5b**), respectively. Of the total new isoforms between 2019 and 2023, 126 and 303 isoforms contained new protein-coding sequence in liver and cerebellar hemisphere, respectively, and 66 and 124 were both medically relevant and have new protein-coding sequence (**Figure 5b**).

**Figure 5:**
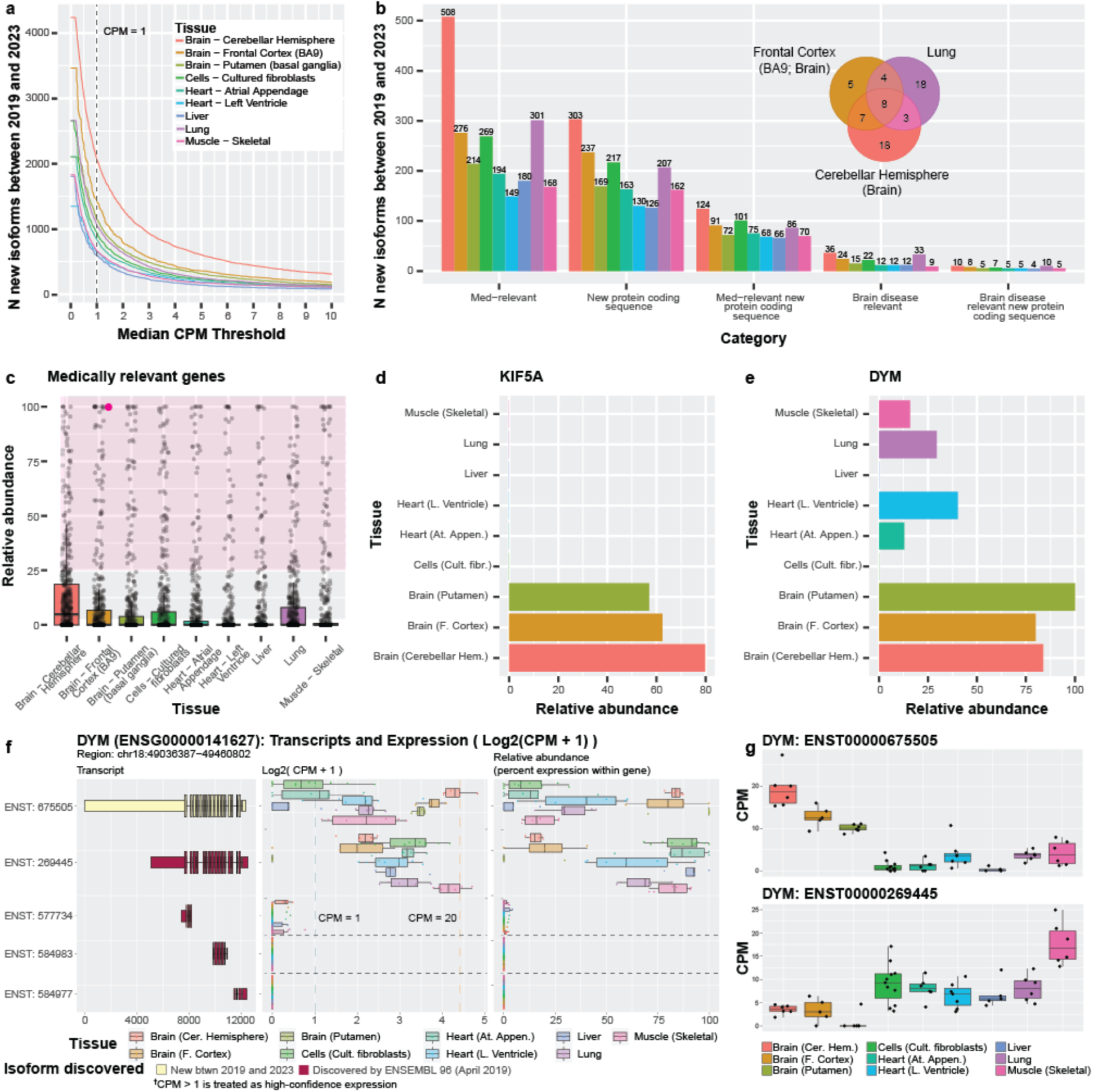
Expression patterns across nine GTEx tissues for isoforms discovered between 2019 and 2023 demonstrate the importance of characterizing and quantifying isoform expression. **(a)** Line graph showing the number of isoforms discovered between 2019 and 2023 expressed across counts-per-million (CPM) thresholds ranging from 0-10. RNA isoforms discovered between 2019 and 2023 expressed in individual tissues above our thresholds (median-unique-counts ≥ 1 and CPM > 1) ranged from 594 (liver) to 2,054 (cerebellar hemisphere). Figures **(b-e)** assume the CPM > 1 threshold. **(b)** Bar graph showing number of isoforms discovered between 2019 and 2023 per tissue from medically and disease-relevant genes. Venn diagram shows overlap of isoforms new between 2019 and 2023 from brain disease relevant genes in cerebellar hemisphere, frontal cortex, and lung. **(c)** Boxplot showing relative abundance of isoforms from 497 medically relevant genes where ≥1 isoforms were annotated between 2019 and 2023, and was expressed in at least one tissue. Each point represents the summed relative abundance for all new isoforms annotated for a given gene discovered between 2019 and 2023. For example, the highlighted point (magenta) represents a gene where its newly annotated isoforms (between 2019 and 2023) constituted 100% of the gene’s expression in frontal cortex. The shaded area highlights genes where its newly discovered isoforms constitute >25% of its total expression for the given tissue. 171 (34.4%) of the 497 genes had a combined relative abundance > 25% in all nine tissues, and new isoforms for 50 (10.1%) constituted >75% of relative abundance in ≥1 tissue. All 497 genes are plotted for each tissue. **(d)** Barplot showing the summed relative abundance of all isoforms new between 2019 and 2023 for *KIF5A*, a brain disease relevant gene. The newly discovered isoforms constitute a major proportion of total expression for the gene. **(e)** Barplot showing relative abundance for the *DYM* isoform discovered between 2019 and 2023 (ENST00000675505), a medically relevant gene. Figures **(f-g)** contain plots from our R-shiny app. **(f)** Plot of five of the isoforms from DYM, showing a cartoon of their exon structure (colored by discovery), the log2(CPM + 1) of the isoforms, and the relative abundance for each isoform. The first isoform was discovered between 2019 and 2023 and is the most highly expressed isoform in several tissues. **(g)** Boxplots show the divergent expression patterns for the top two isoforms from *DYM* suggesting preferential expression between different tissue groups (brain regions vs the rest). Specifically, ENST00000675505 is preferentially expressed in brain while ENST00000269445 is preferentially expressed in the other tissues.

Of the 2,054 new RNA isoforms discovered between 2019 and 2023 that are expressed in the cerebellar hemisphere, 36 are brain-disease relevant, including notable genes such as *MAPT*, *SNCA*, and *KIF5A* (**Figure 5b**). Exactly 33 brain-disease relevant genes are also expressed in lung, which provides an example of disease-relevant genes that may have important functions beyond what they are most associated with. However, most of the brain-disease relevant genes expressed in cerebellar hemisphere and lung do not overlap (Venn diagram in **Figure 5b**). Similarly, 194 of the 829 RNA isoforms discovered between 2019 and 2023 expressed in atrial appendage heart tissue come from medically relevant genes, including *MED12*, *TPM1*, and *DYM*. For lung, 301 of 1053 new isoforms between 2019 and 2023 come from medically relevant genes.

Quantifying all RNA isoform expression patterns for a given gene across tissues is important to understanding its complexity and function, but arguably the “most important” isoforms may be the most highly expressed. Thus, we wanted to assess relative abundance for isoforms annotated between 2019 and 2023 (compared to isoforms already known) in genes where ≥ 1 isoform was discovered in that period. Limiting to only medically relevant gene bodies, we summed the relative abundance for all new isoforms between 2019 and 2023 for each of the 497 genes meeting these criteria. Of these 497 medically relevant genes, the isoforms discovered for 171 (34.4%) genes had a combined relative abundance > 25% in all nine tissues, showing that many of the isoforms discovered in that time period consistently constituted a meaningful proportion of the gene’s expression; this result also highlights the importance of their respective discoveries. Similarly, 50 (10.1%) of the 497 medically relevant genes with isoforms discovered between 2019 and 2023 constituted > 75% relative abundance (**Figure 5c**). For the 452 genes with isoforms containing new protein-coding sequence discovered between 2019 and 2023, the combined relative expression of the new isoforms for 184 (40.7%) of the genes constituted > 25% relative abundance in all nine tissues. Total expression for the new isoforms for 72 (15.9%) of those genes constituted a relative abundance greater than 75% in at least one tissue. (**Supplementary Figure 19; Supplementary Table S8**) *KIF5A* is a brain-disease relevant gene with a large amount of its relative abundance stemming from isoforms discovered between 2019 and 2023, where the four new isoforms comprise 62.4%, 57.0%, and 79.8% of total gene expression for frontal cortex, putamen, and cerebellar hemisphere, respectively (**Figure 5d**). *KIF5A* is implicated in several diseases, including spastic paraplegia 10^60–62^, neonatal intractable myoclonus^63,64^, and amyotrophic lateral sclerosis (ALS)^65–67^. For those with a vested interest in understanding and treating these diseases, knowing about and understanding all of *KIF5A*’s isoforms and their function is essential—perhaps especially in the context of interpreting the functional consequences of genetic variants^2^.

*DYM* is another a medically relevant gene, where mutations are known to cause Dyggve-Melchior-Clausen syndrome (DMC), a type of skeletal dysplasia also known to be associated with brain developmental defects^68,69^. *DYM* was first named in 2003 and has 20 annotated isoforms in Ensembl v109. The overall median gene expression falls between approximately five and 25 CPM for the nine GTEx tissues included here (**Supplementary Figure 20**). Only two isoforms are expressed above our thresholds where one (ENST00000675505) is new between 2019 and 2023, has a new protein-coding sequence, and accounts for most of the gene expression in all three brain regions (frontal cortex: 80.0%; cerebellar hemisphere: 83.9%; putamen: 100.0%; **Figure 5e,f,g**); this isoform is also present in four other tissues (skeletal muscle, lung, left ventricle [heart] and atrial appendage [heart]), but is not the primary isoform expressed. This described expression pattern may imply that these isoforms have tissue-specific roles.

### Newly discovered isoforms from brain frontal cortex are expressed across various tissues

In our recent work by Aguzzoli-Heberle et al.^2^, we discovered a total of 700 new, high-confidence RNA isoforms using deep long-read cDNA sequencing in human prefrontal cortex (Brodmann area 9/46) from twelve aged brain samples (six Alzheimer’s disease cases and six age-matched controls), where 428 of the 700 new isoforms were from known nuclear gene bodies, 267 were from entirely new nuclear gene bodies, and five were spliced isoforms from mitochondrially encoded gene bodies. We directly validated many of these new RNA isoforms from known genes at the protein level via mass spectrometry, along with three from new gene bodies^2^. Observing spliced isoforms from mitochondrially encoded gene bodies was entirely unexpected given dogma dictates that mitochondrially encoded genes are not spliced, but our results supported previous work by Herai et al.^70^ demonstrating the same phenomenon. For interest, we also discovered 2,729 other potential new RNA isoforms^2^, but here, we limit our analyses to those we considered high-confidence (*i.e.,* median CPM > 1 in our previous study). Thus, here, we sought to quantify expression for our high-confidence isoforms across the nine GTEx tissues.

For each tissue we first quantified the proportion of the new isoforms that exceeded our threshold (median unique count ≥ 1 & median CPM > 1; **Figure 6a,d,g**). For new isoforms from known gene bodies (nuclear and mitochondrial), we unsurprisingly found the greatest proportion of isoforms validated in the GTEx frontal cortex samples where 336 (78%) of our recently discovered isoforms were expressed above our required threshold (**Figure 6a**). Of the remaining 92 (22%) that did not meet our threshold, 52 (12%) were observed with 0 < median CPM ≤ 1, while still requiring the median unique counts ≥ 1. The large difference in sequencing depth between the two studies may explain why the remaining 10% did not validate in the GTEx frontal cortex data. Median sequencing depth per sample in Aguzzoli-Heberle et al. was 35.5 million aligned reads per sample^2^, whereas median sequencing depth within the GTEx samples was 4.95 million aligned reads per sample^15^. We also found that 292 (67%) and 300 (69%) of our newly discovered RNA isoforms from known gene bodies were expressed in the cerebellar hemisphere (brain) and putamen (brain), respectively (**Figure 6a**).

**Figure 6:**
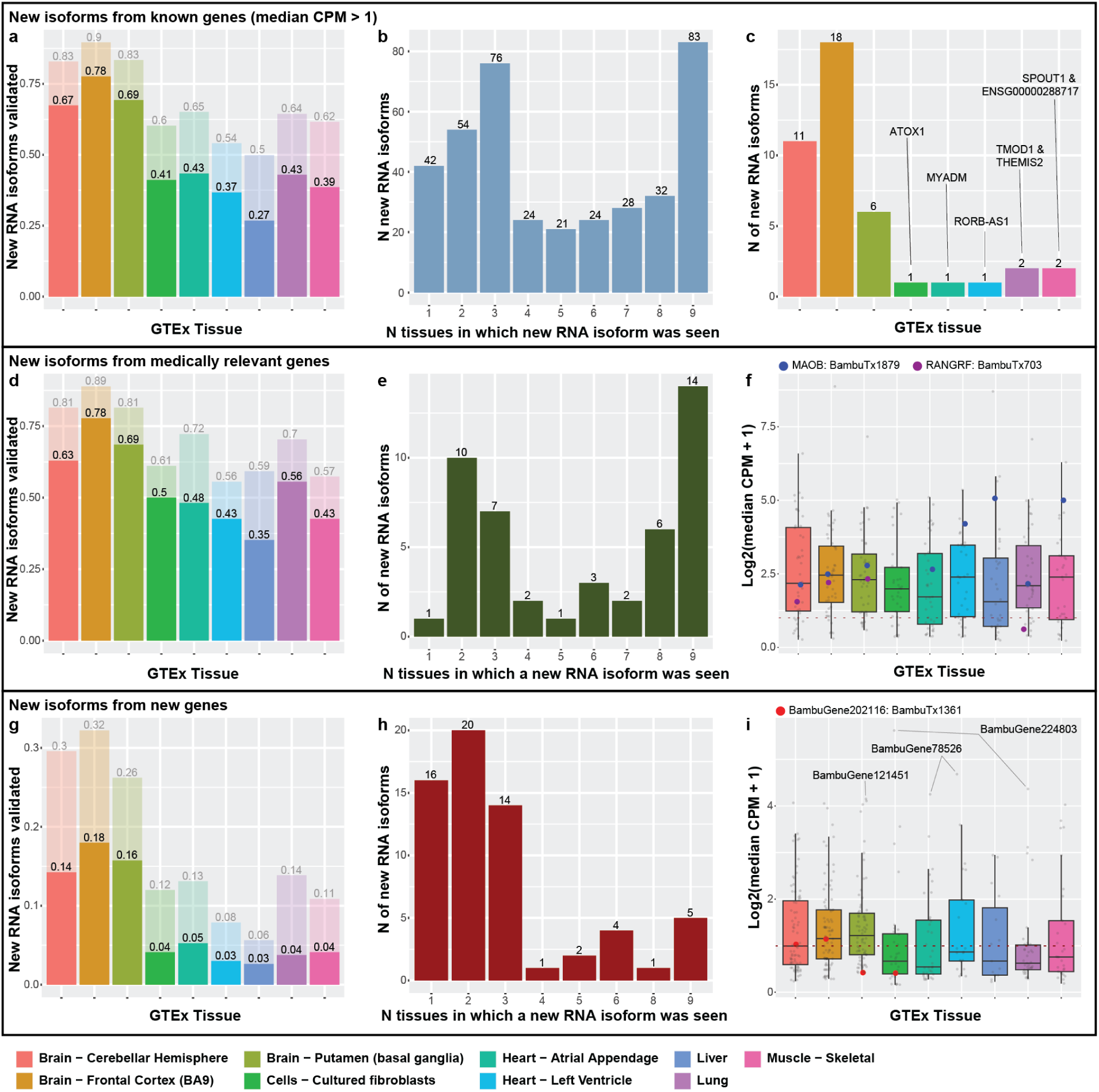
Newly discovered isoforms validated across nine GTEx tissues, including many from new gene bodies. Figures **(a-c)** deal with new isoforms from previously known gene bodies. **(a)** Barplot showing proportion of new isoforms from known genes that validated (at median-unique-counts ≥ 1 and median CPM > 1) for each tissue. Exactly 292 (67%), 336 (78%), and 300 (69%) of our newly discovered RNA isoforms from known gene bodies were expressed in the cerebellar hemisphere, frontal cortex, and putamen, respectively, within the GTEx data. Many new RNA isoforms from known genes were also expressed in the other tissues, ranging from 116 (27%; liver) to 188 (43% atrial appendage). In total, 384 (90%) of the new isoforms from known genes validated in at least one of the nine tissues. **(b)** Barplot of how many tissues in which isoforms were expressed. Of those isoforms that validated in GTEx tissues, 83 were expressed in all nine (largest bar), whereas the next highest group was for isoforms found in three tissues (76). **(c)** Barplot showing tissues where isoforms expressed in only a single tissue were seen. As expected, most isoforms that validated in only a single tissue were found in the prefrontal cortex (18), but several appeared to be “specific” to other, non-brain tissues. Figures **(d-f)** deal with new isoforms from medically relevant genes (subset of new isoforms from known genes). **(d)** Barplot of the proportion of new isoforms from medically relevant genes that validated in each tissue. **(e)** Same as (b) but for medically relevant genes. For figures **(f,i)**, colored dots indicate the expression of isoforms that validated at the protein level in Aguzzoli-Heberle et al.^2^ **(f)** Boxplot of the log2(median CPM +1) of each isoform expressed from the medically relevant genes. Figures **(g-i)** deal with isoforms from new gene bodies. **(g)** Proportion of isoforms from new gene bodies that validated in each respective tissue. **(h)** Same as (b) and (e), but for new gene bodies. **(i)** Boxplot of the log2(median CPM + 1) of each isoform from new gene bodies.

Many of the new RNA isoforms from known genes were expressed in the other tissues, ranging from 116 (27%; liver) to 188 (43% atrial appendage; **Figure 6a**). In total, 447 and 384 isoforms validated in at least one of the nine tissues for all new isoforms and only isoforms from known genes, respectively.

Next, we quantified how many new isoforms from known gene bodies were expressed in multiple tissues (**Figure 6b**). Surprisingly, of those isoforms that validated in the GTEx tissues, 83 were expressed in all nine (largest bar in **Figure 6b**), whereas the next highest group was for isoforms found in three tissues with 76. Observing such a large proportion that validate in three tissues was unsurprising since three of the tissues are from brain—the organ where the isoforms were originally discovered—but observing so many expressed in all nine tissues was surprising. A potential explanation for why such a large proportion validated in all nine tissues could be that many of these RNA isoforms perform essential “housekeeping” functions across tissue types. Only deeper experimental work will be able to determine with certitude.

For the 42 new RNA isoforms from known gene bodies expressed in a single tissue, we assessed which tissue they validated in (**Figure 6b,c**) because isoforms being expressed in a single tissue could indicate tissue “specificity”. Even though these new isoforms were discovered in prefrontal cortex, having greater expression in another tissue may indicate the isoform is “more essential” in that tissue. Unsurprisingly, most of the new isoforms that validated in only a single tissue were found in the prefrontal cortex (18), but several appeared to be “specific” to other, non-brain tissues (**Figure 6c**). In total, there were seven isoforms—one in cultured fibroblasts (*ATOX1*), one in each heart tissue (*MYADM* & *RORB-AS1*), two in skeletal muscle (*SPOUT1* & *ENSG00000288717*), and two in lung (*TMOD1* & *THEMIS2*; **Figure 6c**; **Supplemental Table S9**).

We found similar validation patterns when limiting to new isoforms from known medically relevant genes (**Figure 6d-e**), except only one validated in only a single tissue (*GAP43* in frontal cortex). Median expression for new medically relevant isoforms was comparable across all tissues (**Figure 6f**). All eleven of the new isoforms from known genes that validated at the protein level in our previous work were also expressed in at least one GTEx tissue, two of which were from medically relevant genes (**Figure 6f**).

### Many newly discovered gene bodies are expressed across the nine tissues

In addition to discovering new isoforms from known genes, we also previously discovered 267 isoforms from 245 new gene bodies^2^. Exactly 63 isoforms from the new genes validated in at least one tissue (**Figure 6g,h**), where most that were expressed in only a single tissue were primarily seen in a brain tissue. Only five were expressed in all nine tissues (**Figure 6h**).

Unsurprisingly, most isoforms from the new gene bodies were more highly expressed across the three brain tissues, though the median isoform expression across all tissues was below 3 CPM (**Figure 6i**). Surprisingly, the highest median CPM values for isoforms from new gene bodies did not come from a brain region, however. Cultured fibroblasts and lung expressed BambuGene224803 at median CPM 48.26 and 19.61 respectively. BambuGene78526 has an isoform (BambuTx2076) expressed in Heart - Left Ventricle at 24.71 and Heart - Atrial Appendage at 18.04. The highest median CPM for an isoform from a new gene body in brain came from BambuGene121451, expressed in putamen at 16.78.

Overall, 6.0% (16) of isoforms from new gene bodies were seen in at least one tissue above a CPM of 5. Of the three isoforms from new gene bodies that previously validated via mass spectrometry, only one (1) was expressed in the GTEx samples, and it was only in two of the brain tissues (**Figure 6i**).

### Newly discovered isoforms demonstrate potential tissue specificity or housekeeping roles

We were especially intrigued by the newly discovered RNA isoforms that fell into two specific categories: (1) those that only validated in a single tissue, suggesting potential tissue specificity; and (2) those that validated in all nine tissues, suggesting potential housekeeping roles. In both cases, these newly discovered RNA isoforms could have significant roles in human health and disease. To refine the list of isoforms showing potential tissue specificity or housekeeping roles, we performed differential expression using DESeq2, because simply separating isoforms by the number of tissues in which they are expressed is an overly simplistic approach.

#### Many isoforms exhibit “preferential” expression for specific tissues

Given we have data for two heart regions and three brain regions, we allowed isoforms to be “preferentially expressed” in up to three tissues for this analysis. For our purposes, we designated isoforms as preferentially expressed if they were positively and differentially expressed in up to three tissues compared to all other tissues (pairwise). Specifically, we required a log2-fold change ≥ 1, a false discovery rate (FDR) less than 0.1, and the isoform had to be upregulated relative to the other tissues. We were not interested in isoforms with lower expression relative to the other tissues because this would not suggest tissue specificity. We only included isoforms expressed with a median CPM > 1 in at least one tissue.

We identified 23 isoforms that were preferentially expressed in a single tissue, 46 preferentially expressed in two tissues, and 68 in three tissues (**Figure 7a**). Exactly 22 were from new genes (5 in single tissue, 8 in two tissues, and 9 in three tissues). Unsurprisingly, most of these isoforms were preferentially expressed in at least one brain tissue (12, 39, and 62 from single tissue, two tissues, and three tissues, respectively).

**Figure 7:**
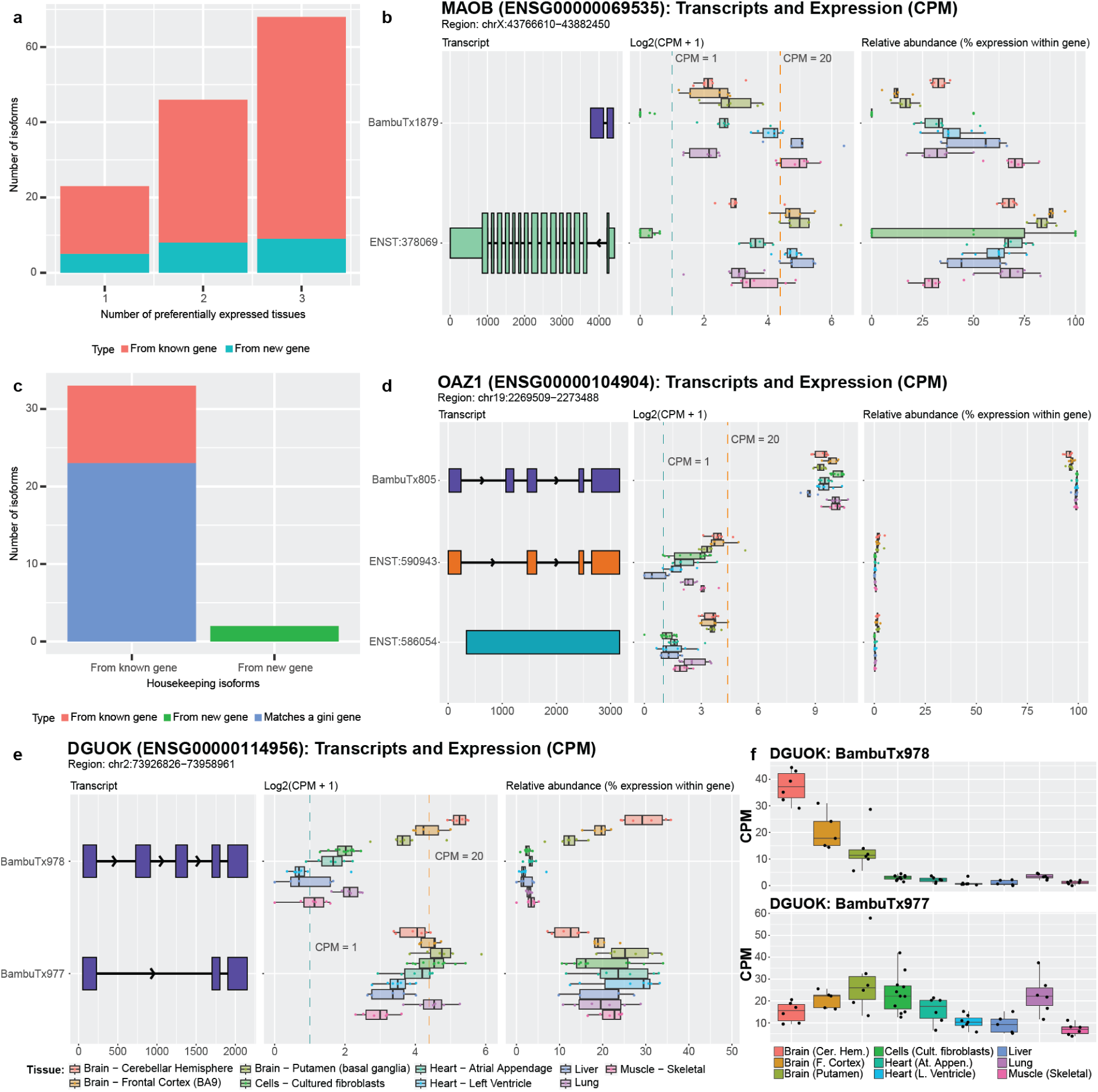
Newly discovered isoforms may have preferential expression and/or housekeeping potential. **(a)** The number of newly discovered isoforms that are preferentially expressed across one, two, and three tissues range from 18 to 59. For new gene bodies, they range from 5 to 9. (b) *MAOB* expression and relative abundance. The new isoform (BambuTx1879) is much shorter than the primary isoform but has a new exon. It is preferentially expressed across liver, muscle, and left ventricle (see Supplemental Figure 22 for unedited plot); whether the new isoform is biologically functional is unknown. (c) The number of newly discovered isoforms that are expressed across all tissues within a log2 fold change of each other for known and new gene bodies, meeting our criteria for housekeeper expression patterns. 23 of the 35 (66%) isoforms that met our criteria for housekeeping expression patterns came from genes that were already identified as housekeeping genes by Joshi et al.^73^, supporting our hypothesis that the other isoforms that met our criteria may also have housekeeping functions, including our newly discovered genes from Aguzzoli-Heberle et al.^2^. (d) *OAZ1* expression and relative abundance. *OAZ1* was previously identified as a housekeeping gene, though its housekeeping status has been debated. The new *OAZ1* isoform is consistent with our defined housekeeping expression pattern, being expressed across all nine tissues and are within log2 fold change of each other (see Supplemental Figure 23 for unedited plot). Whether *OAZ1* fits criteria for a housekeeping gene may vary according to which isoform(s) are being measured in a given assay (*e.g*., different PCR primers). (e) The two newly discovered *DGUOK* isoforms suggest the gene may serve both preferential and housekeeping biological needs. One isoform (BambuTx978) appears to be preferentially expressed, while the other (BambuTx977) exhibits housekeeping-like expression behavior (see Supplemental Figure 24 for unedited plot) (f) CPM values of the two DGUOK isoforms, showing clearly unique expression profiles across the nine tissues included in this study.

Isoforms preferentially expressed in tissues other than brain were especially intriguing, given they were discovered in brain tissue. *MAOB*, for example, encodes an enzyme that helps break down neurotransmitters (often in the brain)^71^ and is a target for treating Parkinson’s disease symptoms. Parkinson’s patients suffer from tremors, rigidity, bradykinesia (slow movement), and resting tremors, in part because of insufficient dopamine levels. MAO-B inhibitors are used to decrease *MAOB* activity and increase dopamine levels, therefore mitigating symptoms in Parkinson’s disease patients^72^. According to our analyses, the newly discovered isoform for *MAOB* is preferentially expressed in muscle, liver, and the left ventricle of the heart, which may imply a distinct function in these tissues (median CPM from 17.37 to 32.48, next highest is 5.87 in brain [putamen]; **Figure 7b**). This preferential expression in other tissues highlights the need for a more in-depth interrogation of *MAOB* isoforms.

#### Newly discovered isoforms exhibit potential housekeeping expression patterns

Opposite of genes and RNA isoforms that demonstrate tissue specificity or preference are genes and RNA isoforms that perform housekeeping functions—genes and isoforms that are broadly required across cells and tissues for proper function. Here, we consider isoforms as demonstrating “housekeeping” expression patterns if they were expressed in all nine tissues above our threshold and demonstrated relatively similar expression across all nine tissues (*i.e.*, were within a log2-fold change of two for all tissues). Using these thresholds, we identified 35 of the newly discovered isoforms that met our criteria for housekeeping expression patterns (**Figure 7c**). As potential validation, 23 of these isoforms matched genes listed in Joshi et al.^73^ as housekeeping genes (or Gini genes) using GTEx data, based on the Gini coefficient—a statistical measure of the inequality among groups, often used in economics where a lower Gini coefficient indicates lower income inequality. We further calculated the Gini coefficient for each of the 35 isoforms and found that 19 isoforms had a Gini coefficient < 0.3; 13 of the isoforms were from Gini genes listed in the above paper (**Supplemental Figure 21**).

One of the Gini genes is *OAZ1*, which has been listed as a housekeeping gene^73,74^, though some publications disagree with this assignment^75,76^. These discrepancies could be due to the lack of annotation for the newly discovered isoform which we classify as housekeeping and is highly expressed (median CPM from 413.71 to 1341.99; **Figure 7d**). In addition, two of the housekeeping isoforms we found come from newly discovered genes.

There were two genes, *DGUOK* and *EEF1AKMT1*, that were especially interesting because they both had an isoform that met our criteria for preferential expression while the other isoform met our criteria for a potential housekeeping isoform. Both genes were also identified as housekeeping genes by Joshi et al.^73^. One of these genes, *DGUOK,* encodes an enzyme that is critical in mitochondria. As most cells contain mitochondria, it is unsurprising that *DGUOK* would be classified as a housekeeping gene based on the Gini index and that a specific isoform could be the primary contributor for its housekeeping status, as is the case with one of the new isoforms, BambuTx977. The other new isoform (BambuTx978), however, is preferentially expressed in brain (**Figure 7e,f**), which may be because the brain has high energy demands and may require tissue-specific functions from *DGUOK.* Based on this hypothesis, however, it is unclear why an additional isoform would be required instead of simply increasing expression for another isoform in brain regions. Only deeper investigation will be able to assess whether these new *DGUOK* isoforms have essential housekeeping and brain-specific functions, respectively.

### Web application for GTEx RNA isoform expression

Using the GTEx data from this study, we created a resource for fellow researchers to explore and query this RNA isoform expression data across nine human tissues. The web app can be found at https://ebbertlab.com/gtex_rna_isoform_seq.html.

## Discussion

Alternative splicing has long been known to play an important role in the biology of complex organisms, like humans, but our understanding of the role various isoforms play is limited. In our opinion, it seems unlikely that 20,000+ protein-coding genes performing a single function can enable such complex organisms^2^. Thus, here, we have provided a broad survey of the RNA isoform landscape to demonstrate both the complexity and diversity of RNA isoforms across nine human tissues using data generated by Glinos et al.^15^ and Aguzzoli-Heberle et al.^2^. We have also tried to demonstrate how little is known about the various isoforms and why it is essential to fully characterize and quantify individual RNA isoform expression across the human cell types, tissues, and lifespan. One of the most important next-steps in biology will be to determine the function for individual RNA isoforms for every gene.

Short-read sequencing technologies have been a major boon for assessing total gene expression across a range of tissue and cell types, and across various diseases, but short-read sequencing struggles to accurately quantify expression for individual isoforms. Essentially, expression for every isoform within a gene is collapsed into a single expression measurement because of the technical limitations of short-read sequencing data.

Long-read sequencing, however, provides a major improvement in our ability to accurately characterize and quantify individual RNA isoforms, but here we also demonstrate that accurately quantifying expression for individual RNA isoforms is not a solved problem, as have others^2,13,16^. For example, these studies highlight the challenges associated with quantifying RNA isoforms given that a large percentage of reads cannot be uniquely assigned to a single isoform, though it is still a major improvement over short-read sequencing data.

Perhaps the most important discussion item arising from this work, however, is whether the many isoforms expressed for certain genes have distinctly different, biologically meaningful functions, or are simply biological redundancy or spurious splicing. We already know of various examples where the distinct isoforms for a given gene are not only biologically meaningful, but essential. *BCL-X* (*BCL2L1*)^7^ is a classic example, where one isoform is pro-apoptotic (BCL-Xs) while the other is anti-apoptotic (BCL-XL). Other examples (*e.g.*, *RAP1GDS1* and *TRPM3*) are known to have more subtle functional differences. Subtle differences do not necessarily imply they are insignificant, however. As we continue to learn more about individual isoforms, we expect to find some proportion that are a result of biological redundancy or spurious alternative splicing, but also expect that many will have biological significance. Ultimately, only deeper studies– including experimental work–will be able to fully address these questions.

## Conclusion

As sequencing technology (including preserving natural RNA) and associated algorithms advance, our ability to study individual RNA isoforms will improve dramatically. Here, we provided a broad survey of the RNA isoform landscape, demonstrating the isoform diversity across nine tissues and emphasizing the need to better understand how individual isoforms from a single gene body contribute to human health and disease. We found genes whose isoform expression patterns differed in interesting and potentially significant ways and we validated isoforms recently discovered in Aguzzoli-Heberle et al.^2^. We also identified isoforms that exhibit patterns consistent with preferential expression for a given set of tissues, and others that demonstrated potential housekeeping expression patterns. The breadth and depth of what can and should be studied is vast and warrants significant efforts if we are to understand the complexities and subtleties of human health and disease.

## Methods

### Downloading and Comparing Ensembl annotations

Gene transfer format (GTF) files for ten representative Ensembl GRCh38 annotations spanning ten years (2014-2023; one per year) were downloaded from the Ensembl website (**Supplemental Table S2**). Using Python version 3.11.3 within a Jupyter notebook (version 6.5.4)^77^ (CODE AVAILABILITY), we quantified the number of genes and isoforms per year, as well as the number of protein-coding genes based on annotations within the GTF file (**Supplemental Table S2**).

Specifically, for a gene or transcript (i.e., isoform) to be considered protein-coding, their respective “gene_biotype” and “transcript_biotype” must have been “protein_coding”. We performed set comparisons between the years 2019, 2021, and 2023 to identify the overlap between annotations. We used the Ensembl v109 (2023) annotations to quantify the most common transcript biotypes, as shown in **Figure 2e**. To calculate the number of annotated isoforms per gene and per protein-coding gene (shown in **Figure 2f,g**), respectively, we summed the number of unique transcripts (based on Ensembl transcript ID) for each unique Ensembl Gene ID. We also calculated the percentiles for annotated isoforms per gene using R version 4.3.1. Annotations from 2014 and 2015 had additional alternative contigs that we excluded from these analyses.

### Downloading GTEx data, read pre-processing, genomic alignment, and quality control

We obtained the publicly available GTEx nanopore long-read cDNA RNAseq data from Glinos et al.^15^ for this study through the AnVIL portal^78^. The data consists of 88 GTEx samples from 15 different human tissues and cell-lines. The data were re-processed using the same methods used in our recent paper by Aguzzoli-Heberle et al.^2^.

Briefly, we pre-processed the cDNA data with Pychopper (version 2.7.6), applying settings compatible with the PCS109 sequencing kit, since the GTEx data were sequenced using this chemistry. Pychopper discards any reads missing primer sequences on either end and recovers reads that contain primer sequence in the middle, which result from fused molecules. Pychopper also orients the reads according to their genomic strand and removes any adapter or primer sequences. We then aligned the pre-processed reads to GRCh38 with minimap2 version 2.26-r1175^79^, including the “-x splice” (to allow spliced alignments) and “-uf” (to identify splicing sites using the transcript strand) alignment parameters. We used the GRCh38 reference genome without alternate contigs for alignment. We removed reads that mapped with a Mapping Quality (MAPQ) score below 10 using samtools (version 1.17). The BAM files were then sorted by genomic coordinates and indexed using samtools. See **Code Availability** for all scripts. Due to minor differences between our pipelines (*i.e.*, we employed Pychopper), the number of reads we included for isoform quantification in Bambu is lower than what Glinos et al. used. See **Supplemental Table S1** for number of reads analyzed by Bambu. See Glinos et al.^15^ for quality control data for these samples.

### Sample inclusion criteria

Glinos et al.^15^ originally sequenced 88 cDNA GTEx samples across 15 tissues and cell lines using the Oxford Nanopore Technologies MinION. For our analyses, we ultimately only included nine of the 15 tissues after applying the following inclusion criteria: (1) we only included tissues with samples from at least five unique subjects; (2) excluded samples with experimental conditions (i.e., *PTBP1* knockdown); (3) excluded technical replicates; (4) excluded samples with <1,000,000 reads; and (5) excluded any samples that did not cluster with their respective tissue group based on a principal component analysis (PCA). In the end, we retained 58 samples across nine tissues, including cerebellar hemisphere (brain), frontal cortex (brain), putamen (brain), cultured fibroblasts, atrial appendage (heart), left ventricle (heart), liver, lung, and skeletal muscle. When selecting which sample to retain among technical replicates, we chose the sample with the highest number of total reads that was below the maximum total reads for that tissue (to avoid including a sample that would be an outlier). We then performed a PCA analysis, using the DESeq2^80^ “plotPCA” function after DESeq2 normalization on the total counts matrix from Bambu (excluding the filtered samples) and excluded one liver sample due to poor clustering (**Supplemental Figures S1,2**). The full list of included samples can be found in **Supplemental Table S1**.

### GTEx analyses

To limit false positives and to be consistent with our recent work in Aguzzoli-Heberle et al.^2^, we selected CPM = 1 as the noise threshold and further required a median-unique-counts ≥1, to exclude any isoforms that have not been consistently observed uniquely across samples. Unless otherwise specified, we only included isoforms with a median CPM > 1 (and median-unique-counts ≥ 1) in our analyses. We calculated counts-per-million (CPM), gene CPM, relative abundance, and the median relative abundance for each isoform using Bambu’s “total counts” metric. For reference, relative abundance is the percent expression for a given isoform within a gene. We used Ensembl v109 annotations (2023) combined with the 700 new isoforms discovered in Aguzzoli-Heberle et al.^2^ to quantify isoform expression with Bambu. We quantified the number of isoforms expressed within each tissue across a range of CPM thresholds from 0-10.01 in 0.01 increments, as shown in **Figure 3a** and further stratified these analyses across a range of gene and isoform types, including protein-coding isoforms, medically relevant isoforms, protein-coding isoforms from medically relevant genes, and brain disease relevant isoforms. Medically relevant genes were as defined by Wagner et al.^20^ with additions from Aguzzoli-Heberle et al.^2^. We generated an upset plot evaluating the overlap of isoform expression across several combinations of different tissues in rank order using R package ggupset version 0.3.0^81^. We also assessed the significance of protein-coding to non-protein-coding isoforms between cerebellar hemisphere and heart’s left ventricle by running a Chi-square test.

#### Isoform heatmap across nine GTEx tissues and pathway analysis

We wanted to identify genes that have a high number of isoforms expressed, so we created a list of genes that express > 5 isoforms at least one tissue (above our ‘noise’ thresholds described above). Using those genes and the number of isoforms expressed by tissue, we created a clustered heatmap with the raw isoform numbers (not expression values). We performed pathway analyses using Metascape^51^ (accessed January 2024). We also plotted the relationship between total gene expression and number of isoforms expressed and ran a Spearman correlation for the significance of the correlation for the left ventricle.

#### New between 2019 and 2023 isoforms

To assess the isoforms that are new between the Ensembl v96 annotation (released April 2019) and the Ensembl v109 annotation (released February 2023), we retained all Ensembl transcript ID’s from Ensembl v109 that were not present in v96. To quantify the percent of total expression for new isoforms since 2019 (as shown in **Figure 5c**), we summed the relative abundance for those isoforms for a given gene.

### New isoforms from Aguzzoli-Heberle et al.^2^

To assess expression patterns for the 700 new isoforms reported in Aguzzoli-Heberle et al.^2^, we separated the isoforms into new isoforms from known genes (433 isoforms) and new isoforms from new gene bodies (267 isoforms) and created an additional subset of new isoforms from medically relevant genes (54 isoforms). Note that in Aguzzoli-Heberle et al., they reported 53 isoforms in this category—the discrepancy between these numbers is due to filtering out the spliced mitochondrial isoform, which we do not exclude here. We then calculated the proportion of new isoforms present in each tissue at a median-unique-counts ≥ 1 and median CPM > 1 (as well as median CPM > 0) by taking the number of isoforms expressed in the group and dividing it by the total number of isoforms possible in the group. We also calculated the number of tissues each isoform was expressed in at a median CPM > 1. Taking the isoforms that were only expressed in a single tissue, we looked at which tissue was expressing the isoform. We plotted the log2(median CPM + 1) of the isoforms in the new from medically relevant genes and new genes categories at a CPM > 0.

### Potential preferential and housekeeping isoforms from new isoforms

To assess whether any of the new isoforms were potentially preferentially expressed in a given set of tissues, or exhibit housekeeping-like behavior (expressed across all tissues), we performed pairwise differential expression analyses between each tissue pair for each isoform using DESeq2 (normalizations using the total counts matrix with all isoforms, then filtering to only newly discovered isoforms for comparisons). To meet our criteria for preferential tissue expression, we required a log2-fold change > 1, a false discovery rate (FDR) less than 0.1, and the isoform had to be upregulated relative to the other tissues. To meet potential housekeeper criteria, we required isoforms to be expressed in all nine tissues above our noise threshold and demonstrate similar expression across all nine tissues (*i.e.*, within a log2-fold change of two for all pairwise tissue comparisons).

Further, we calculated the Gini coefficient for those isoforms that were discovered in Aguzzoli-Heberle et al.^2^. We did this using the R package DescTools. We chose to use the cutoff of 0.3 to determine the Gini isoforms and then compared them to the isoforms we identified using our pairwise differential expression method and the Gini genes identified in Joshi et al.^73^.

### Figures and tables

We created figures and tables using a variety of python (version 3.11.3) and R (version 4.3.1) scripts. Some figures were downloaded from our Rshiny app (R version 4.3.0). All scripts are available on GitHub (see code availability). Isoform structures were visualized using R package ggtranscript^82^ (version 0.99.9). Final figures were assembled using Adobe Illustrator.

### Data availability

GTEx long-read RNAseq data used is available through the AnVIL project^78^ at the following link: https://anvil.terra.bio/%23workspaces/anvil-datastorage/AnVIL_GTEx_V9_hg38. All resulting data from this work will be uploaded to the GTEx AnVIL project for proper data management. Data can be visualized on the Ebbert Lab website at https://ebbertlab.com/gtex_rna_isoform_seq.html and the deep long-read frontal cortex data from Aguzzoli-Heberle can be viewed at https://ebbertlab.com/brain_rna_isoform_seq.html.

### Code Availability

All code used in the manuscript is available at: https://github.com/UK-SBCoA-EbbertLab/landscape_of_RNA_isoform_expression

## Supporting information

Supplemental figures and table descriptions

## Acknowledgments

We appreciate the contributions of the Sanders-Brown Center on Aging at the University of Kentucky. We are deeply grateful to the research participants and their families who make this research possible. We would like to thank the University of Kentucky Center for Computational Sciences and Information Technology Services Research Computing for their support and use of the Morgan Compute Cluster and associated research computing resources. We would like to thank Singularity Sylabs for providing support and extra cloud storage for our software containers.

