## Supplemental figures and table descriptions for "Surveying the landscape of RNA isoform diversity and expression across 9 GTEx tissues using long-read sequencing data"

Supplementary Tables:

Supplementary Table 1: Samples used by tissue with the number of primary aligned reads with MAPQ >= 10 from our analysis

Supplementary Table 2: Ten Ensembl annotations (one per year) used as an overview from 2014-2023 and their download paths.

Supplementary Table 3: Gene and transcripts that were labeled as protein-coding in 2019 and 2023 but not in 2021

Supplementary Table 4: Number of isoforms expressed per gene, based on a unique counts threshold (across all samples combined; unique counts thresholds—1, 5, 10, and 20)

Supplementary Table 5: Number of genes expressing at least N isoforms (based on unique counts, all samples combined) from all genes, protein-coding genes, and medically relevant genes.

Supplementary Table 6: Number of genes expressing at least N isoforms (by tissue) from all genes or protein-coding genes.

Supplementary Table 7: Genes expressing more than five isoforms in at least one tissue and the number of isoforms expressed per tissue for those genes.

Supplementary Table 8: Number of genes expressing isoforms new since 2019 at different relative abundance thresholds.

Supplementary Table 9: List of new isoforms (from Aguzzoli-Heberle et al.) expressed in a single tissue above our noise thresholds.

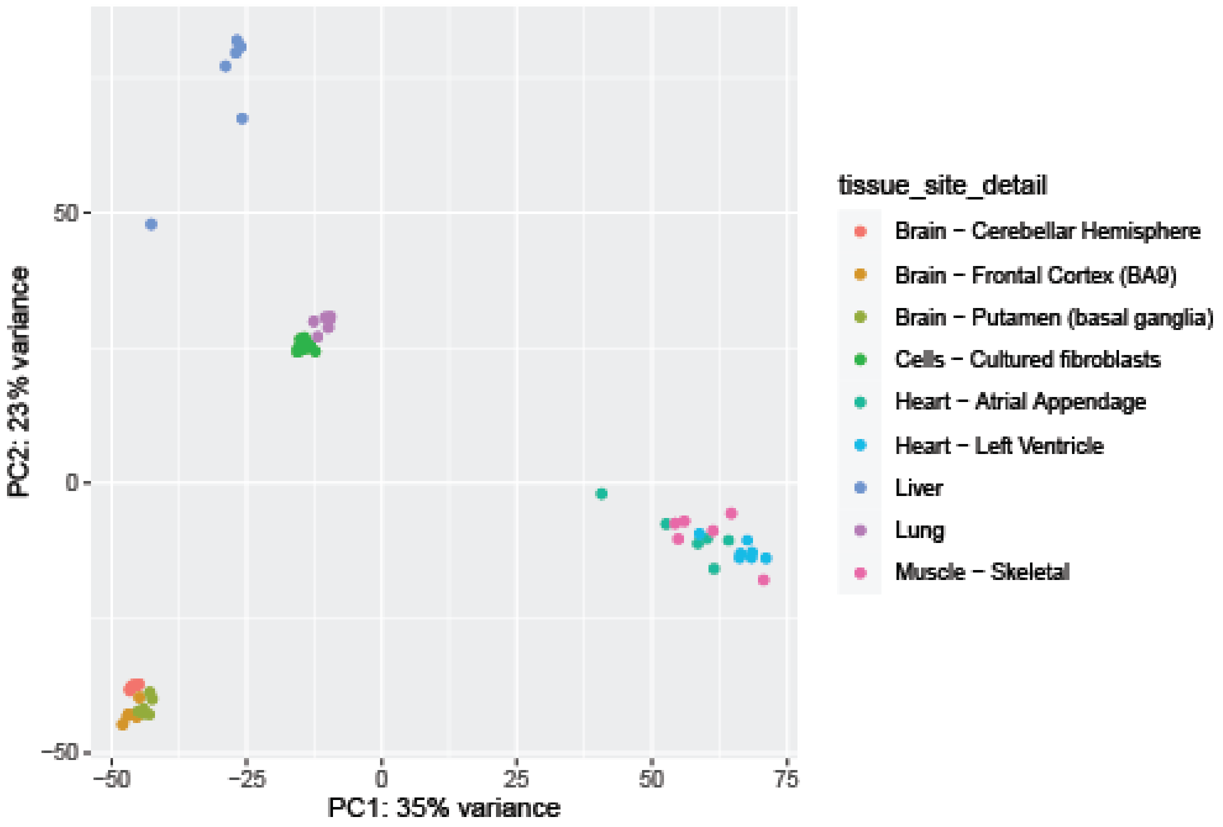


*Supplementary Figure 1: DESeq2 PCA after filtering out technical replicates/experimental samples and those with low read counts.*

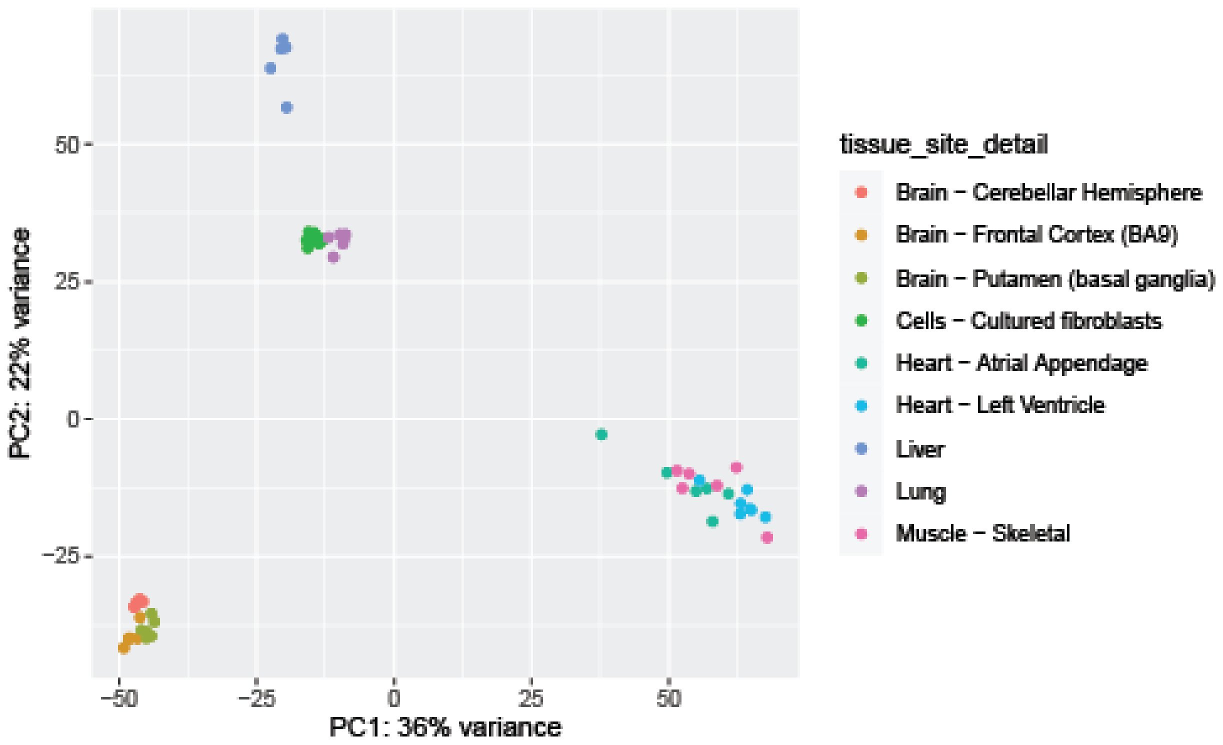


*Supplementary Figure 2: Final DESeq2 PCA after sample with low clustering is removed.*

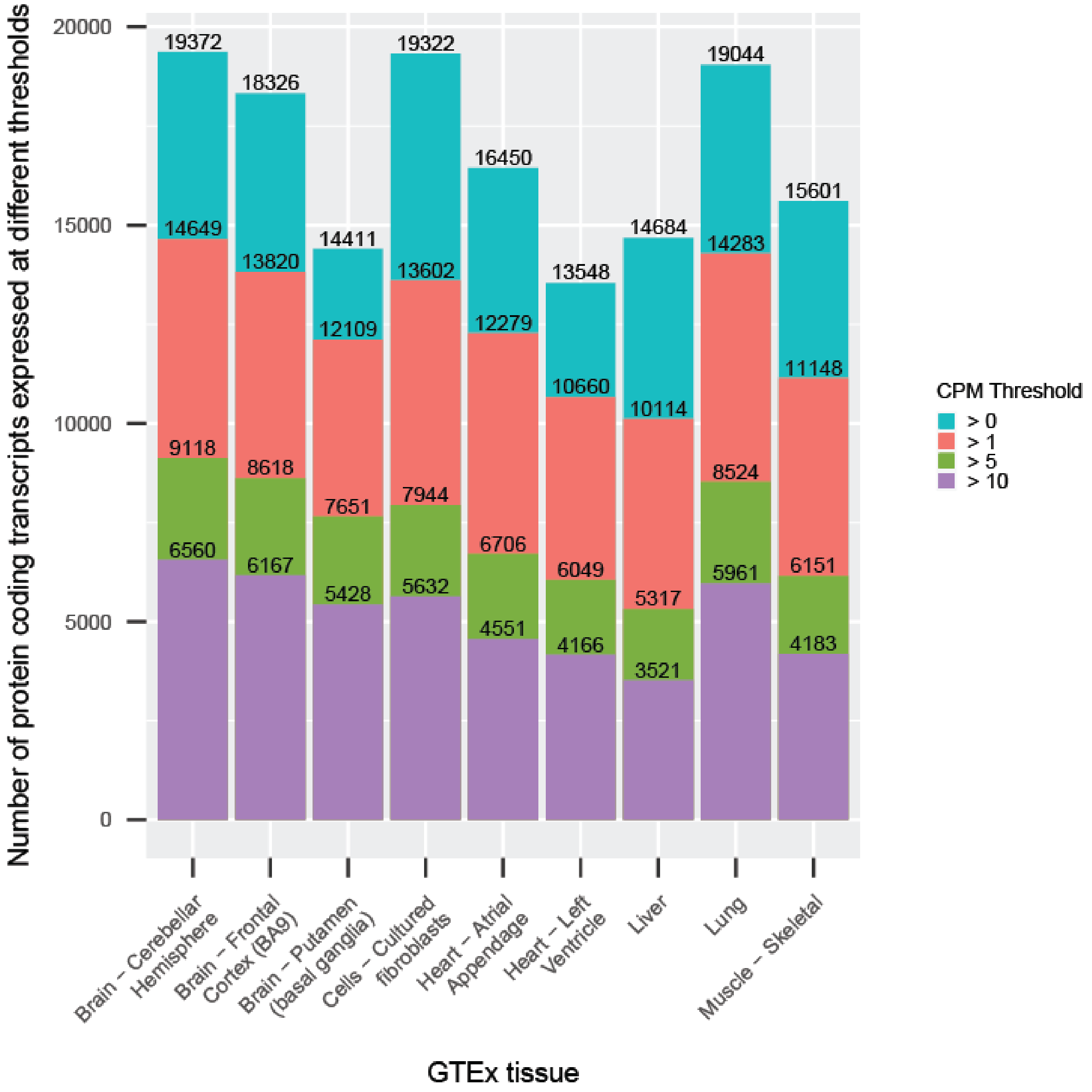


*Supplementary Figure 3: Number of protein-coding isoforms expressed in each tissue across four thresholds.* Number of protein-coding isoforms that passed four different thresholds in each tissue. The pink is the normal threshold used throughout this paper. Other's are reported for interest.

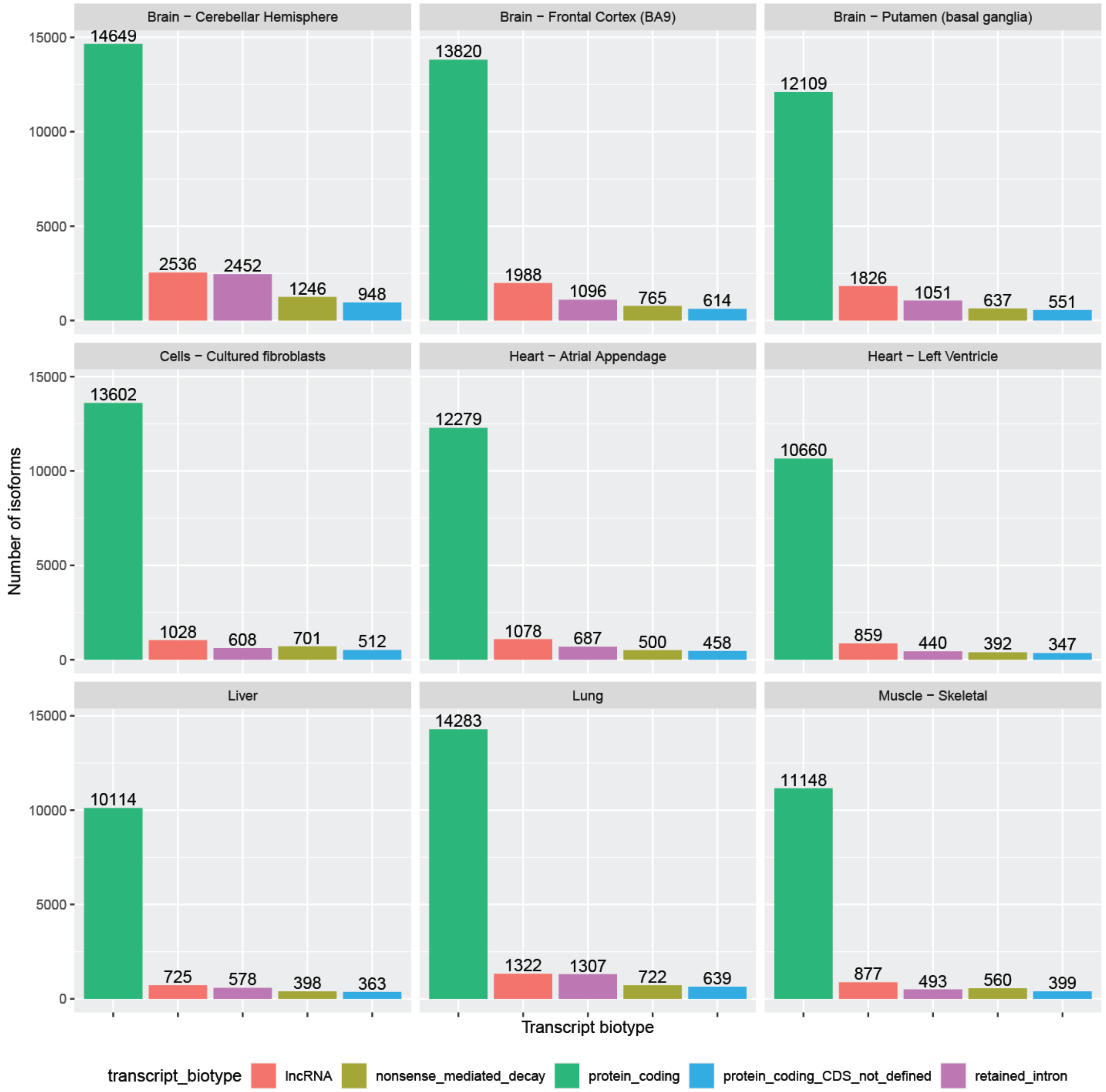


*Supplementary Figure 4: Top five transcript biotypes expressed for each tissue.* When compared with <fig2e> the order of the most expressed biotypes does not match the order of the most annotated transcript biotypes. Note, skeletal muscle and cultured fibroblasts also differ from the other tissues in that nonsense mediated decay is a more prevalent biotype than retained intron.

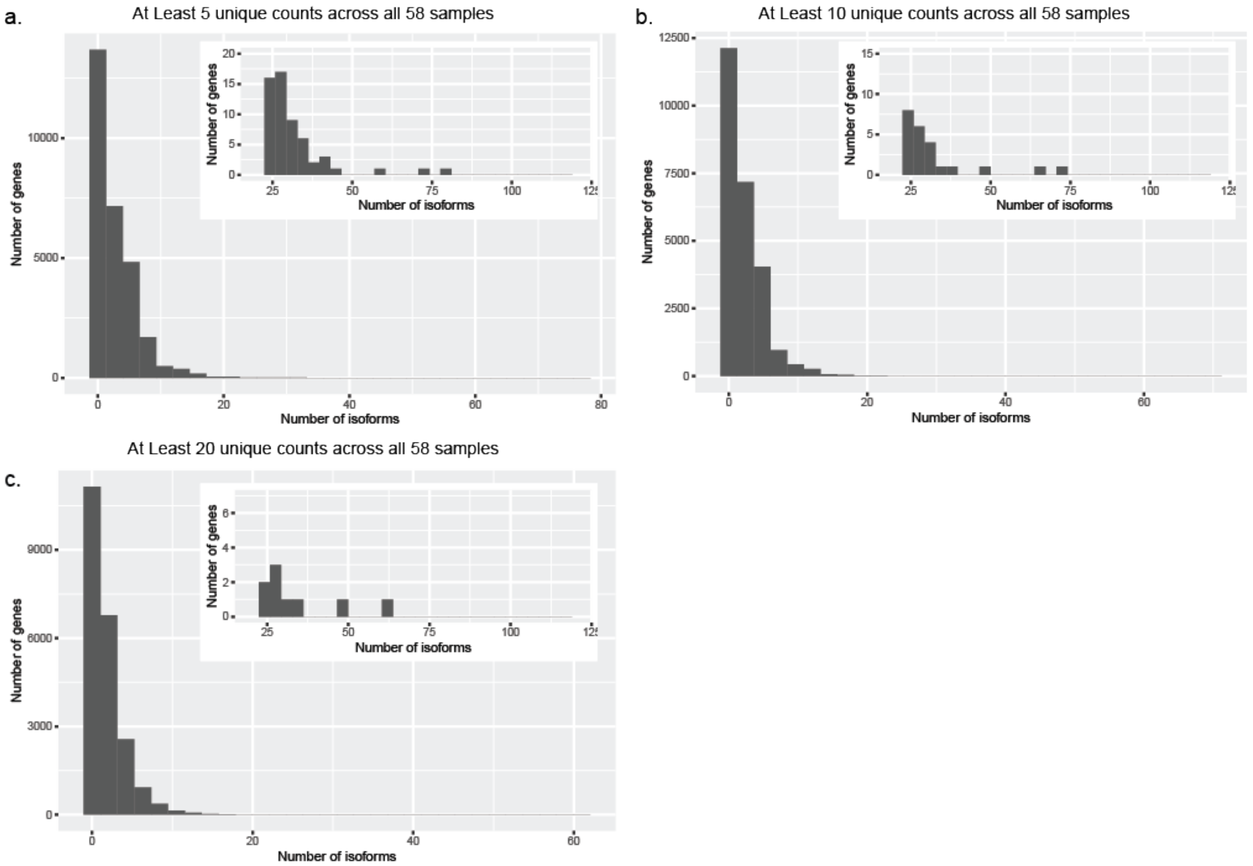


*Supplementary Figure 5: Isoforms expressed per gene based on unique counts.* Histograms of number of isoforms expressed per gene with at least **a.** 5, **b.** 10, or **c.** 20 unique counts across all 58 samples. Nested plots show a zoomed in portion of the histogram.

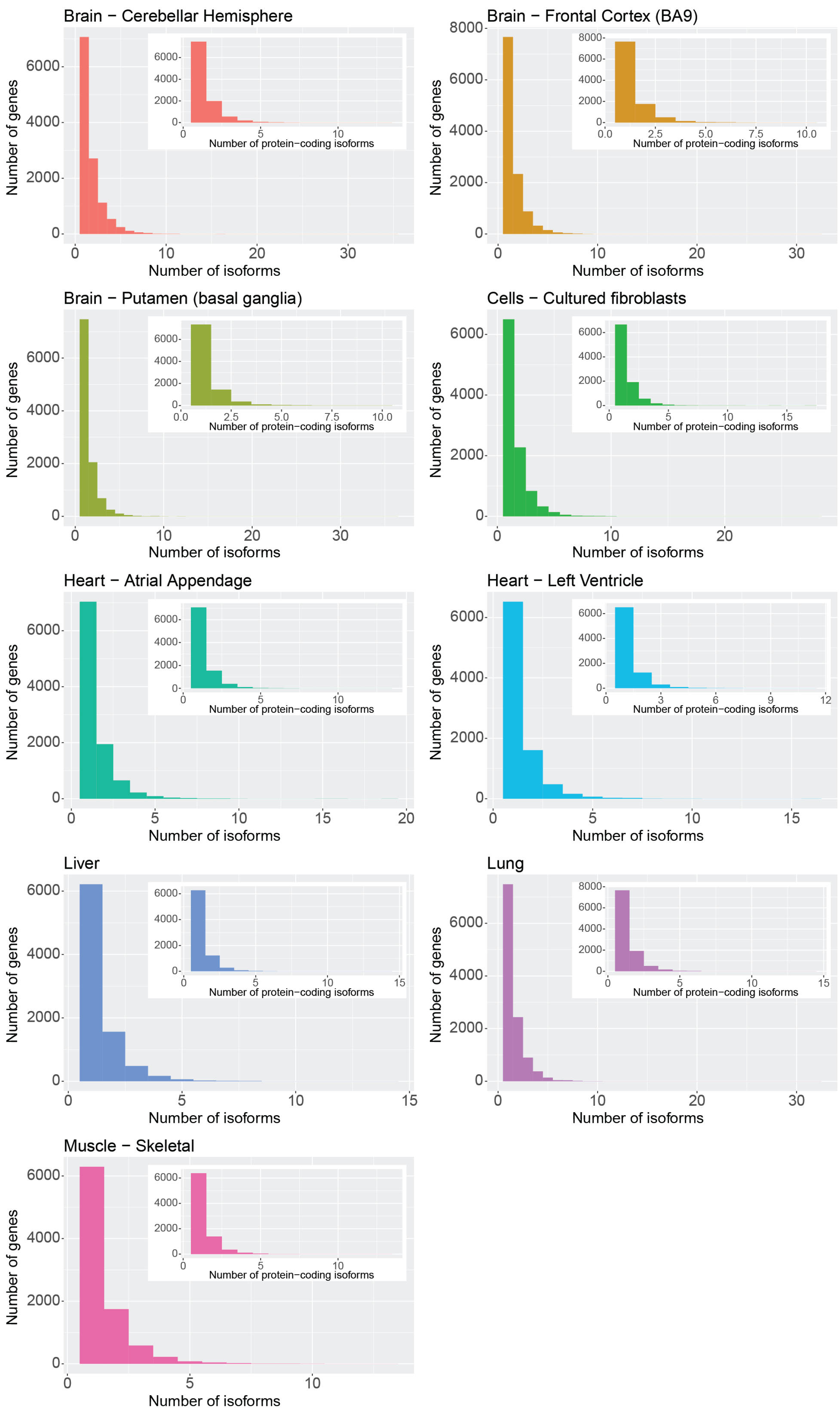


*Supplementary Figure 6: Isoforms expressed per gene by tissue.* Histograms of number of isoforms expressed per gene. Nested histograms show the number of protein-coding isoforms expressed per gene.

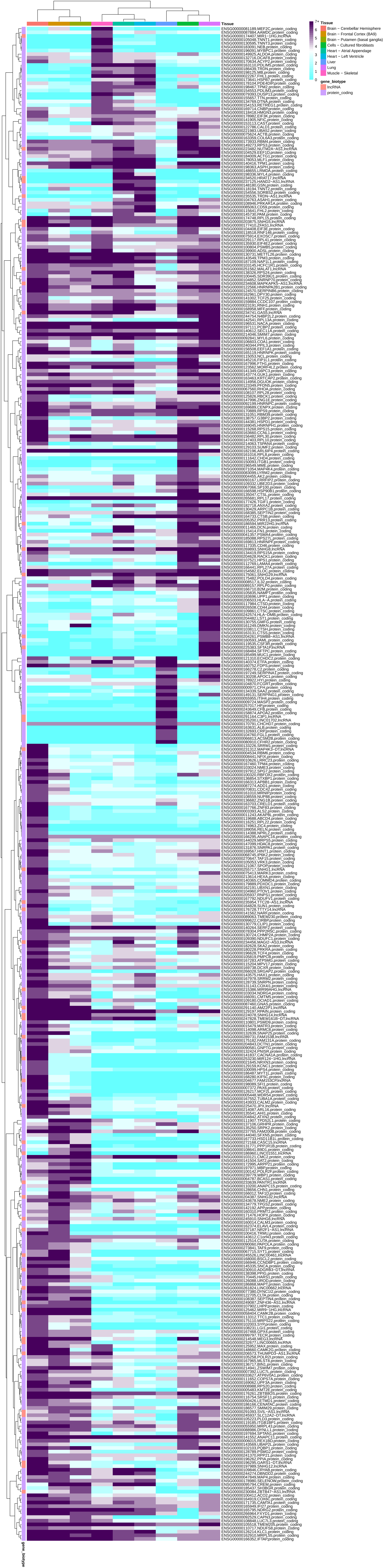
7+
6
5
4
3
2
1
0
Tissue
Brain − Cerebellar Hemisphere
Brain − Frontal Cortex (BA9)
Brain − Putamen (basal ganglia)
Cells − Cultured fibroblasts
Heart − Atrial Appendage
Heart − Left Ventricle
Liver
Lung
Muscle − Skeletal
gene_biotype
lncRNA
protein_coding

*Supplementary Figure 7: Clustered heatmap of number of isoforms expressed per gene.* Clustered heatmap of 416 genes that expressed more than five isoforms in at least one tissue, including the GeneID's, Gene symbol (if available), and gene biotype (if available).

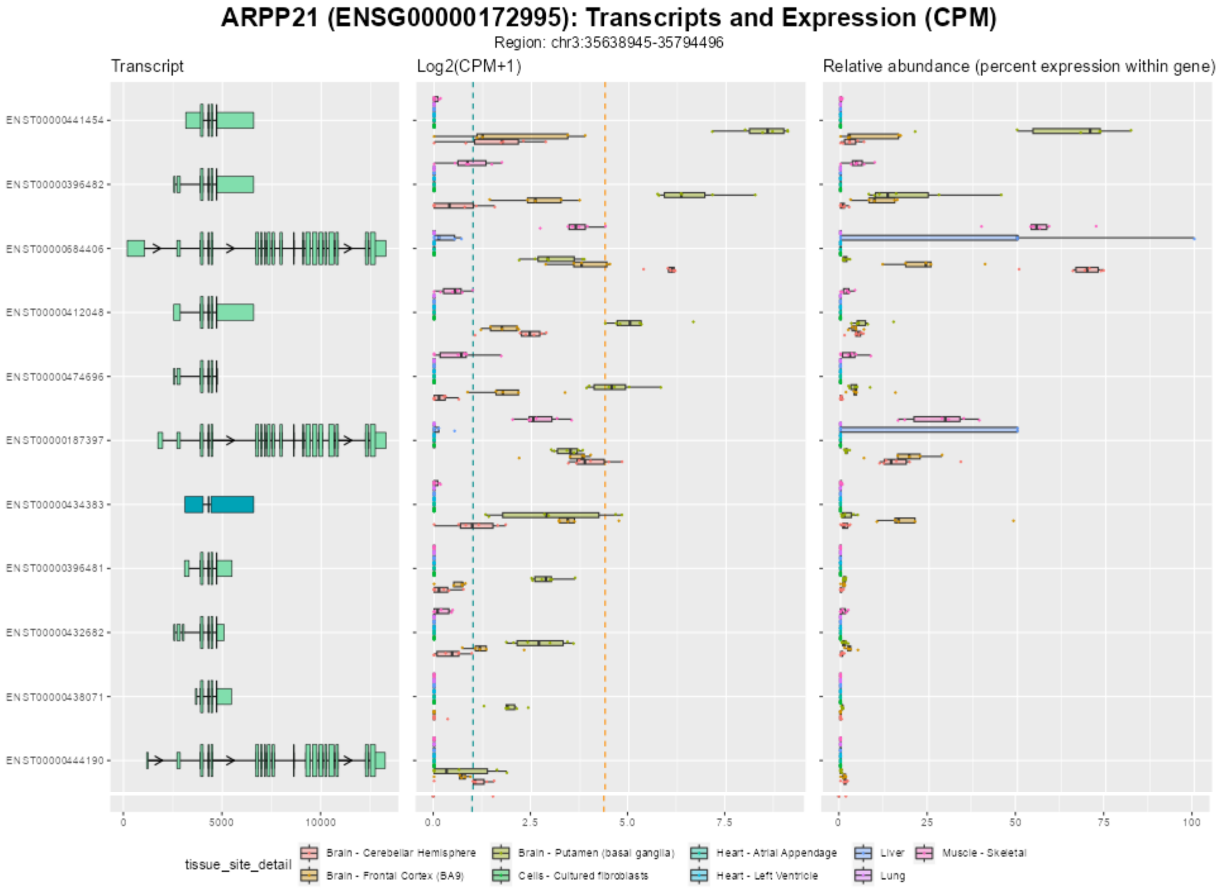


*Supplementary Figure 8: ARPP21 isoforms, expression, and relative abundance.* Isoforms expressed above a median CPM > 1. Putamen expressing the greatest number of isoforms.

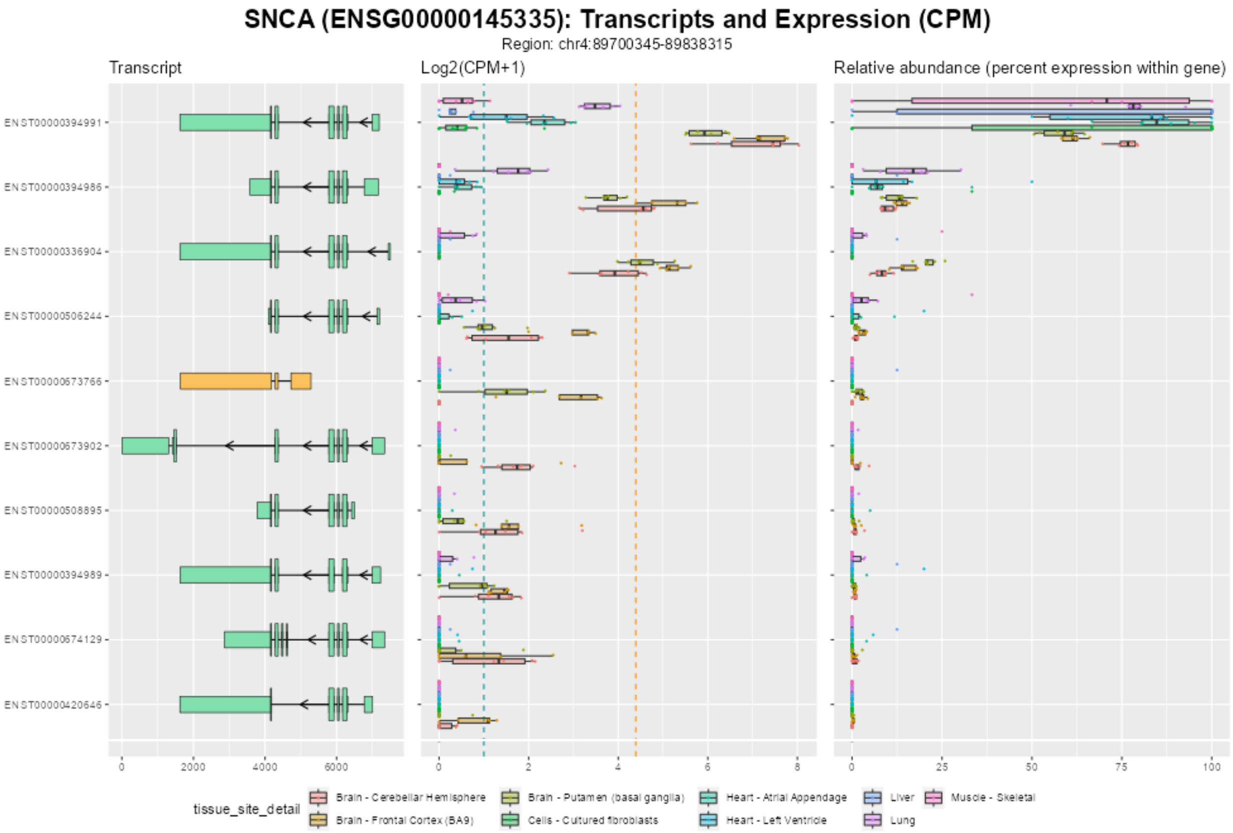


*Supplementary Figure 9: SNCA isoforms, expression, and relative expression.* Isoforms expressed above a median CPM > 1.

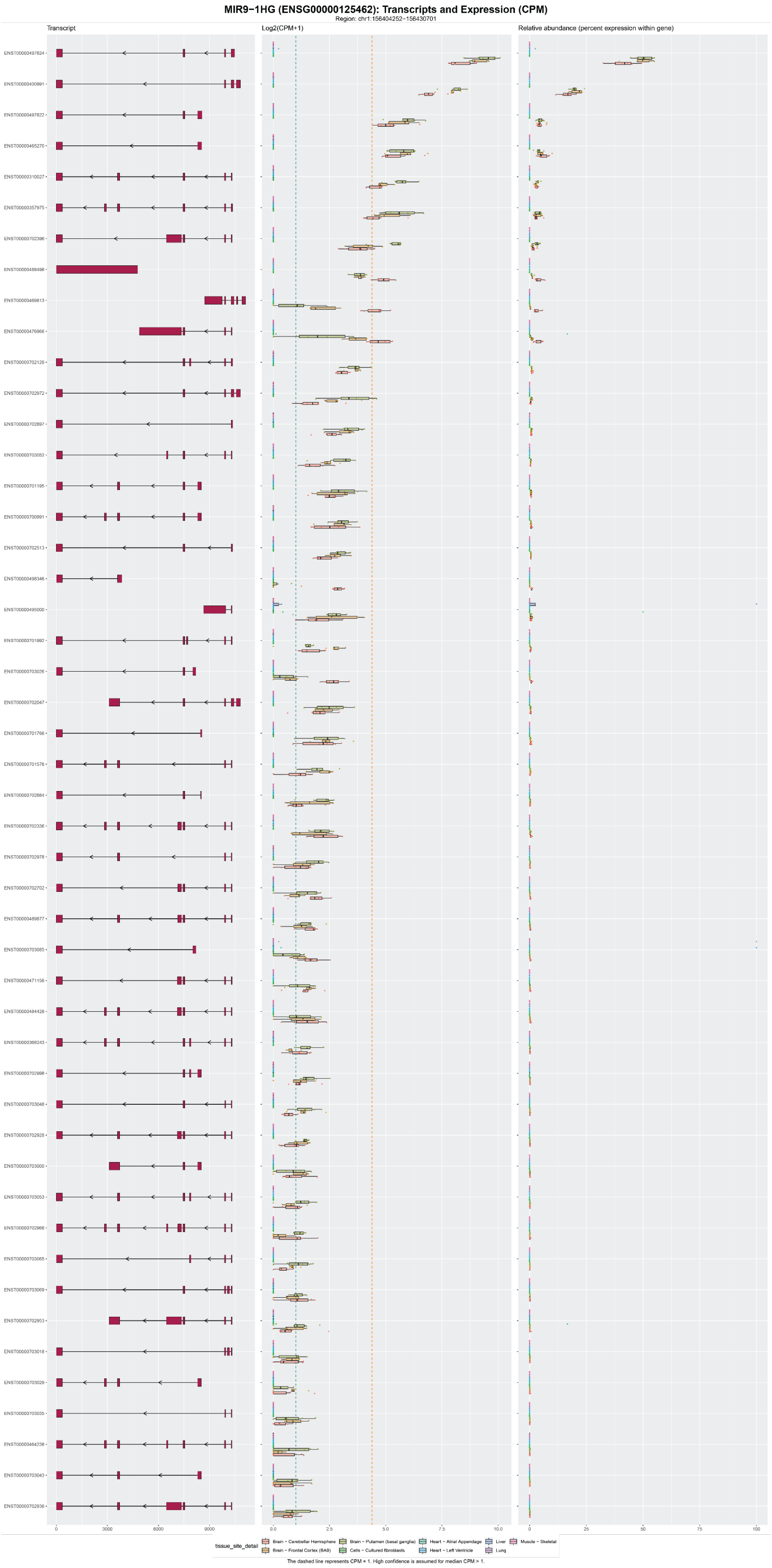


*Supplementary Figure 10: MIR9-1HG isoforms, expression, and relative expression*

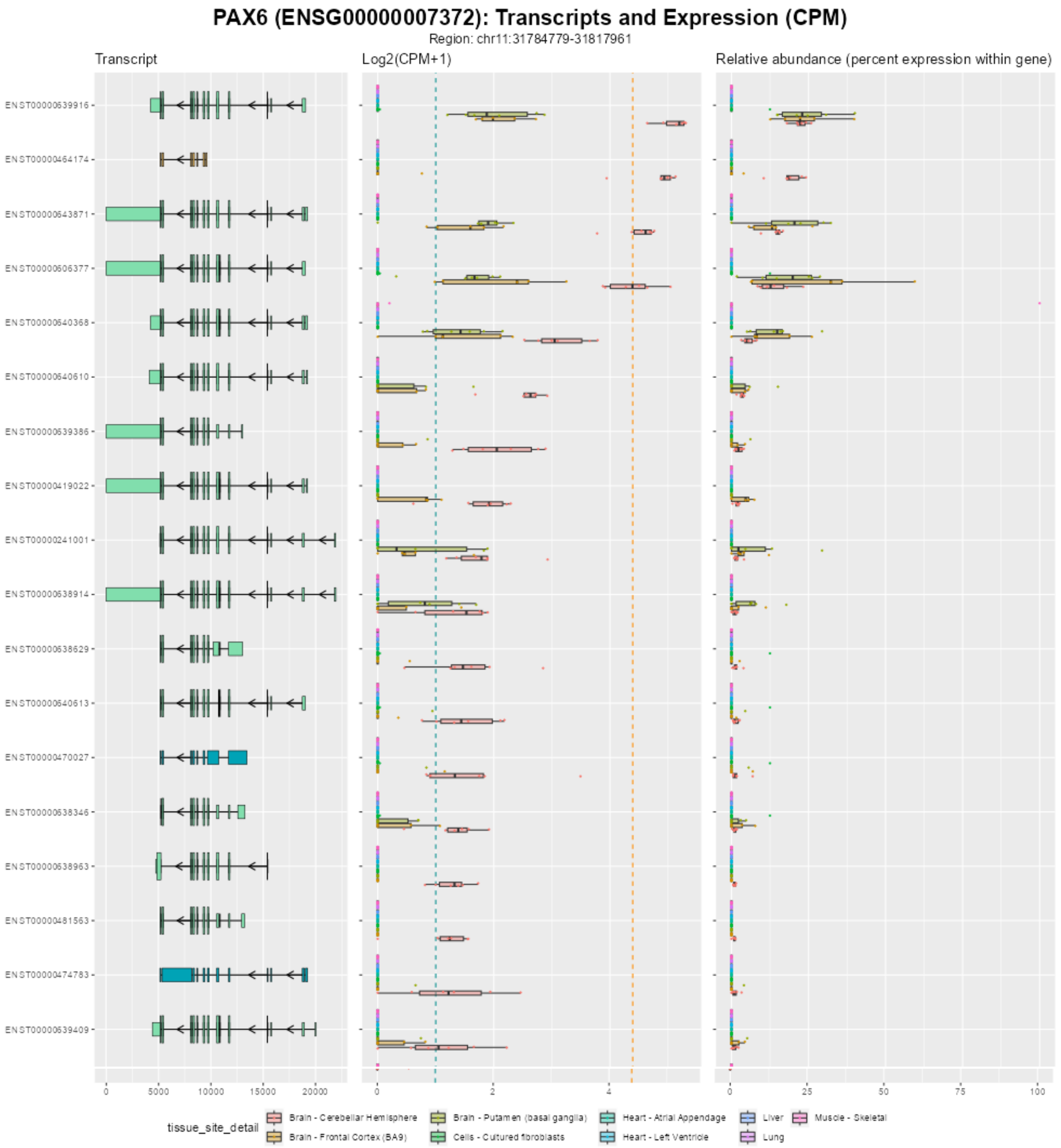


*Supplementary Figure 11: PAX6 isoforms, expression, and relative abundance.* Isoforms only expressed at a median CPM > 1. Relative abundance for each isoform seems similar across the brain tissues.

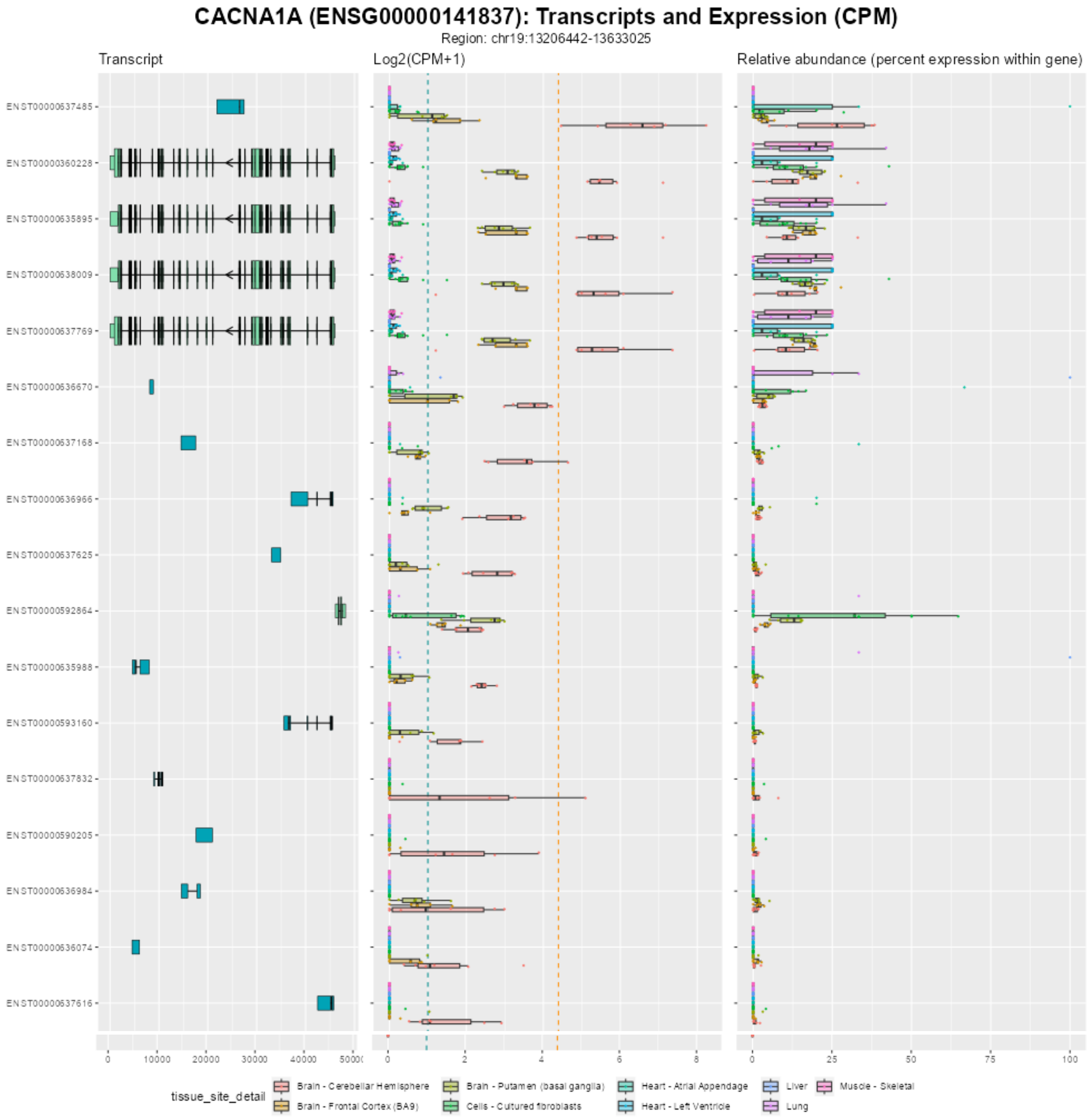


*Supplementary Figure 12: CACNA1A isoforms, expression, and relative expression.* Isoforms expressed above median CPM > 1. Cerebellar hemisphere is expressing the greatest number of isoforms. Relative abundance of isoforms is fairly similar across the top four protein-coding isoforms (seafoam green isoforms).

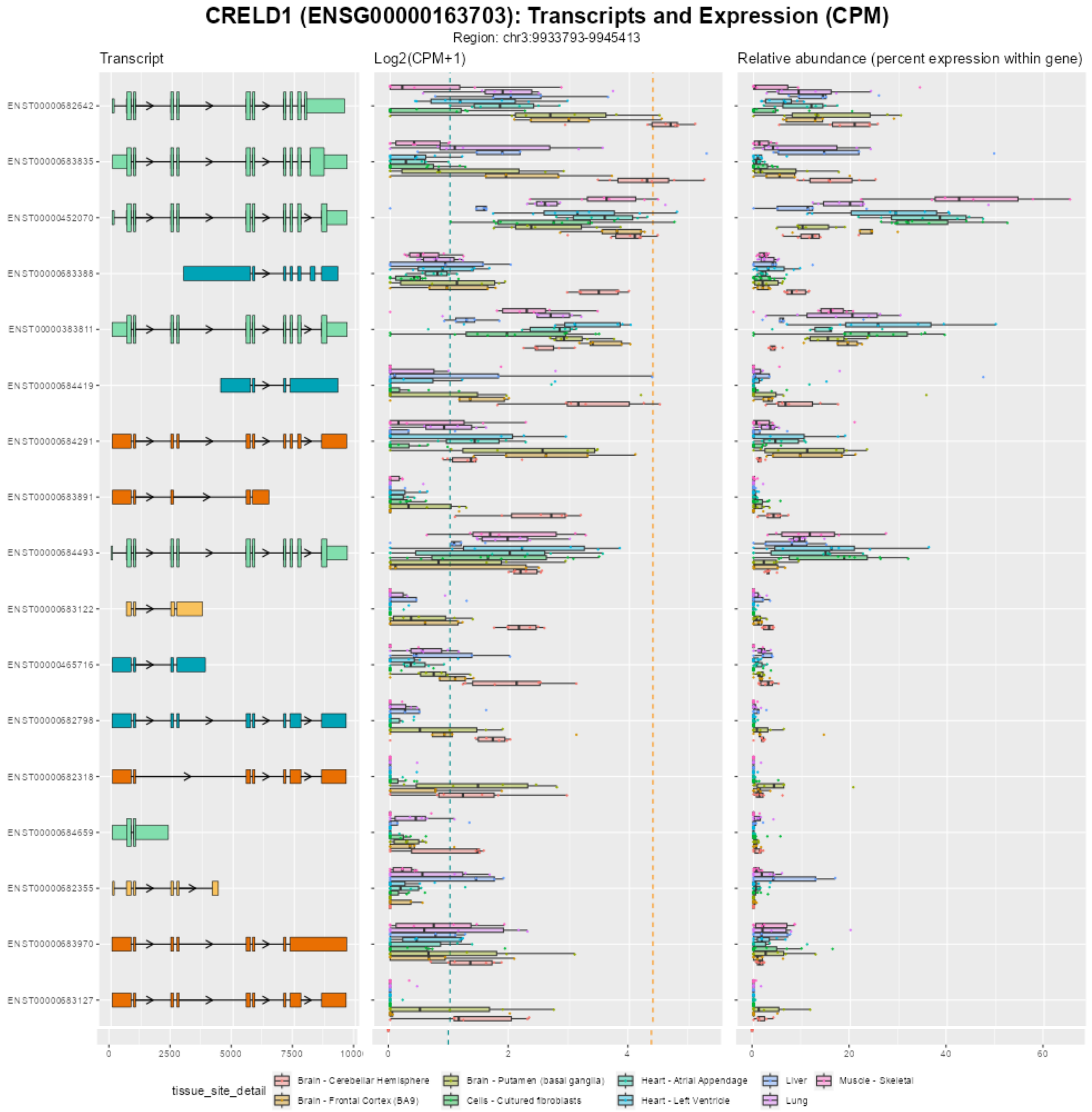


*Supplementary Figure 13: CRELD1 isoforms, expression, and relative expression.* Isoforms expressed above a median CPM > 1. Cerebellar hemisphere expresses the greatest number of isoforms.

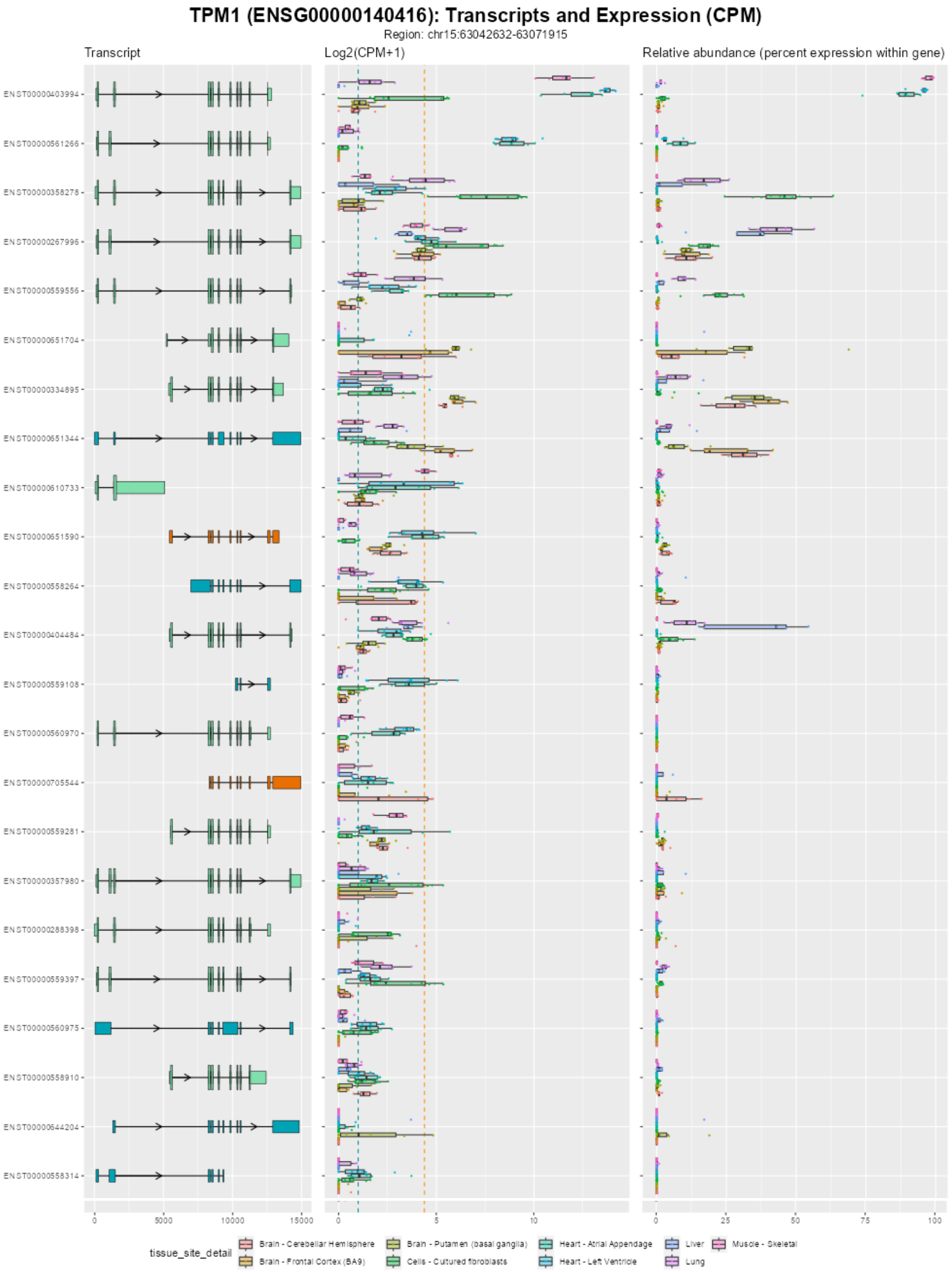


*Supplementary Figure 14: TPM1 isoforms, expression, and relative expression.*

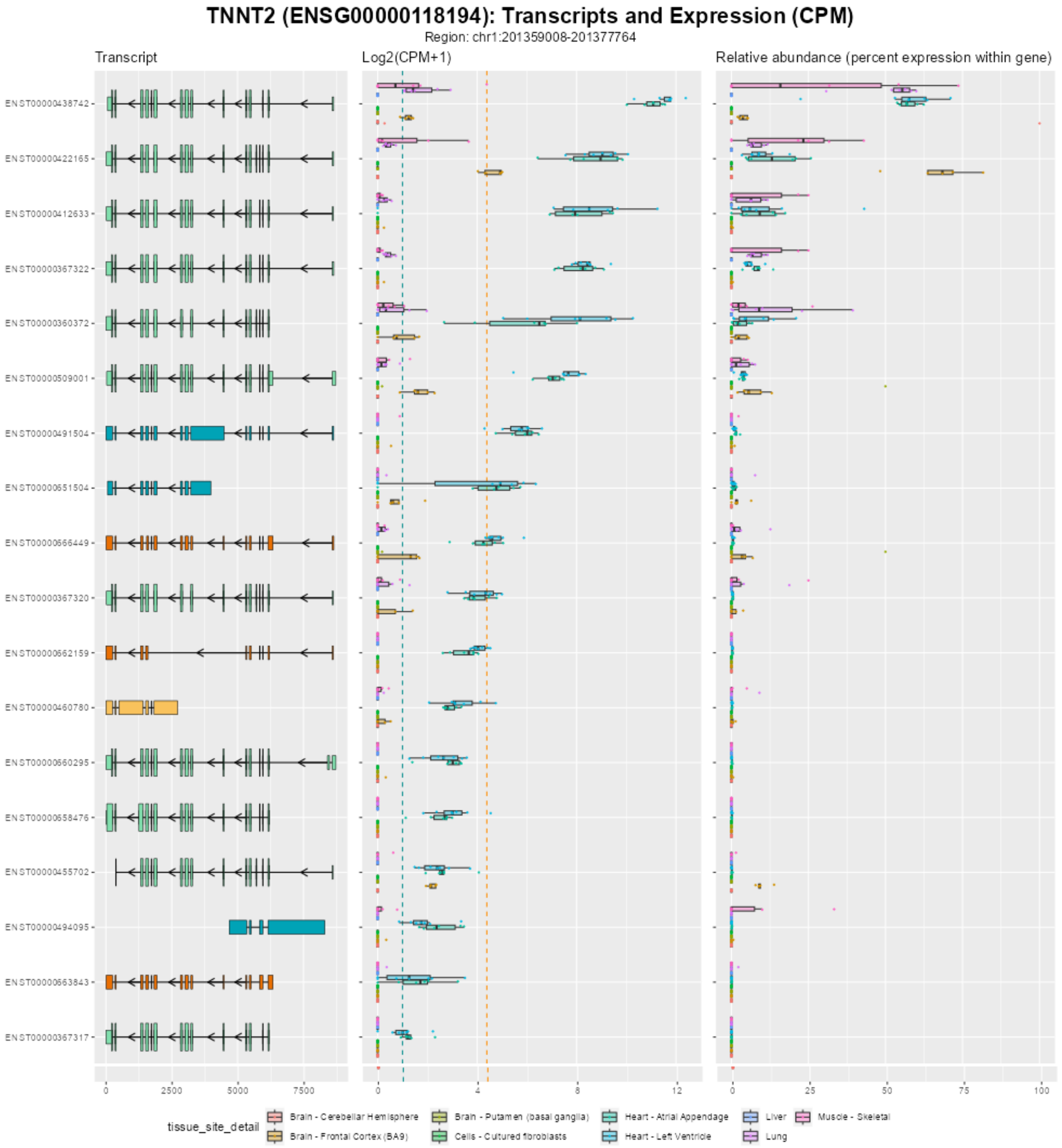


*Supplementary Figure 15: TNNT2 isoforms, expression, and relative expression*

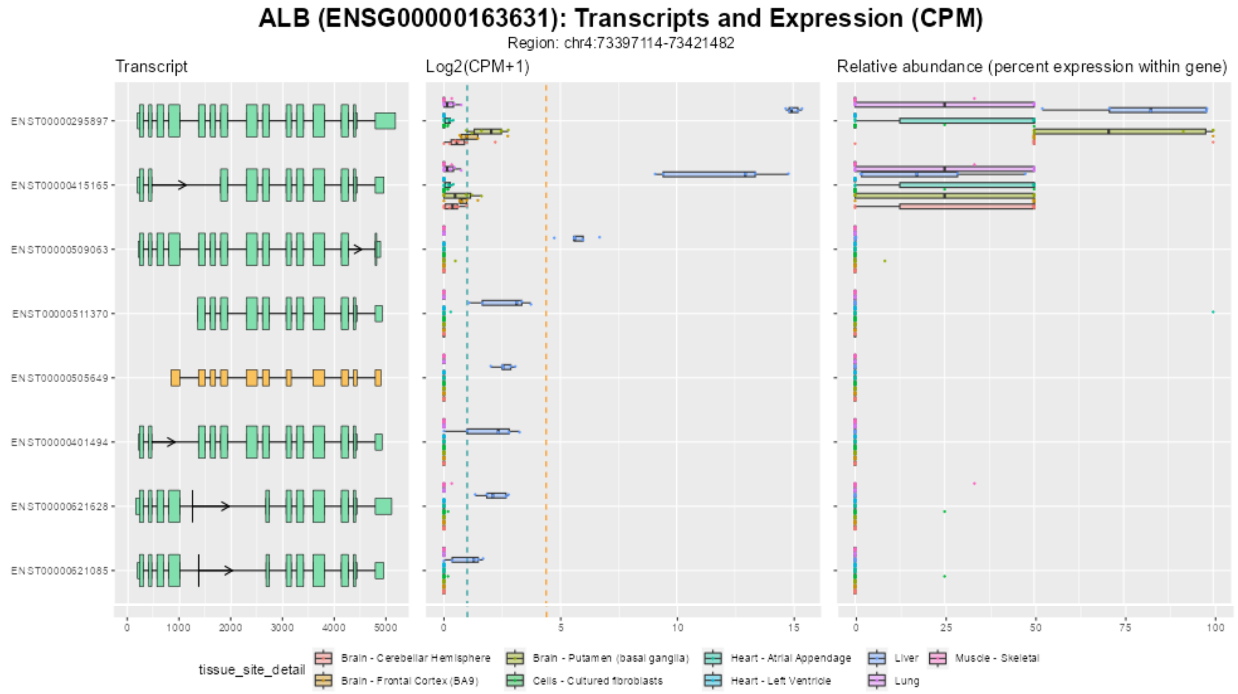


*Supplementary Figure 16: ALB isoforms, expression, and relative expression*

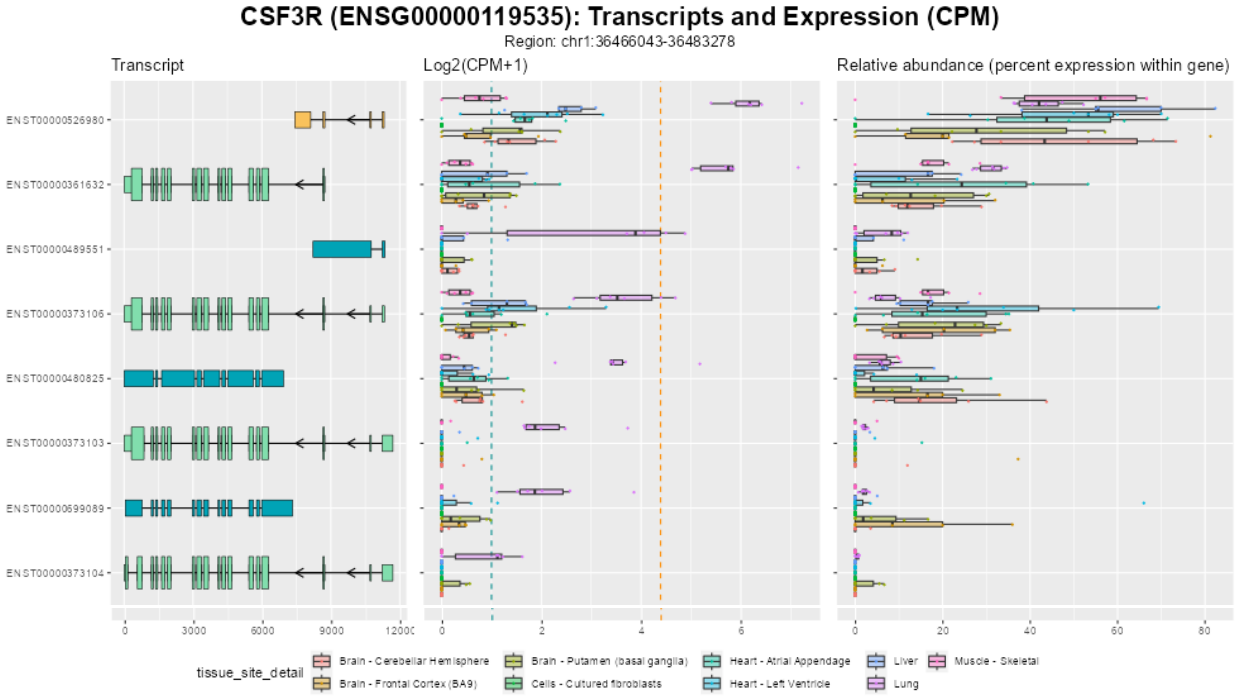


*Supplementary Figure 17: CSF3R isoforms, expression, and relative expression*

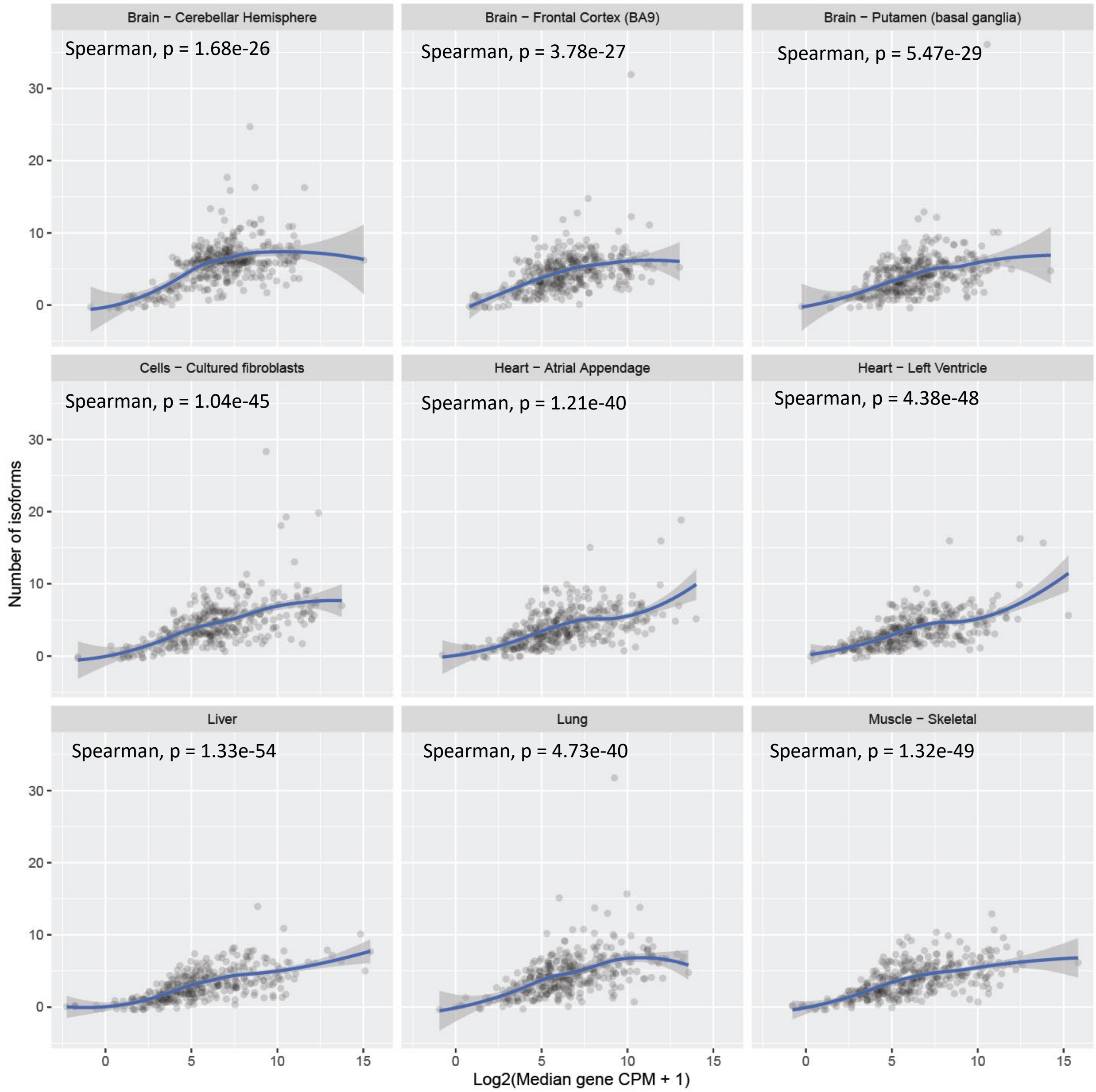


*Supplementary Figure 18: Number of isoforms verses gene expression*

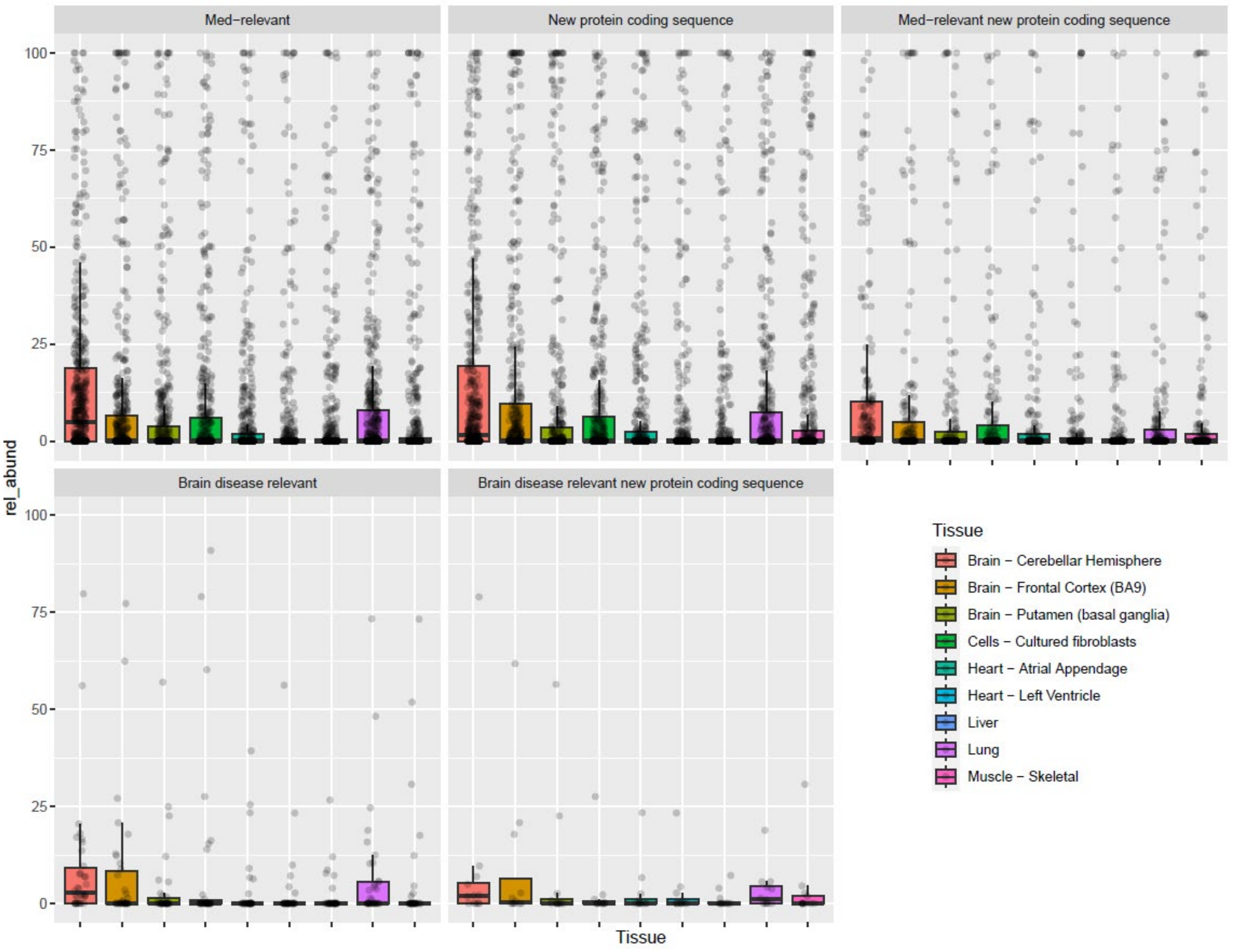


*Supplementary Figure 19: Percent relative abundance across isoform categories*

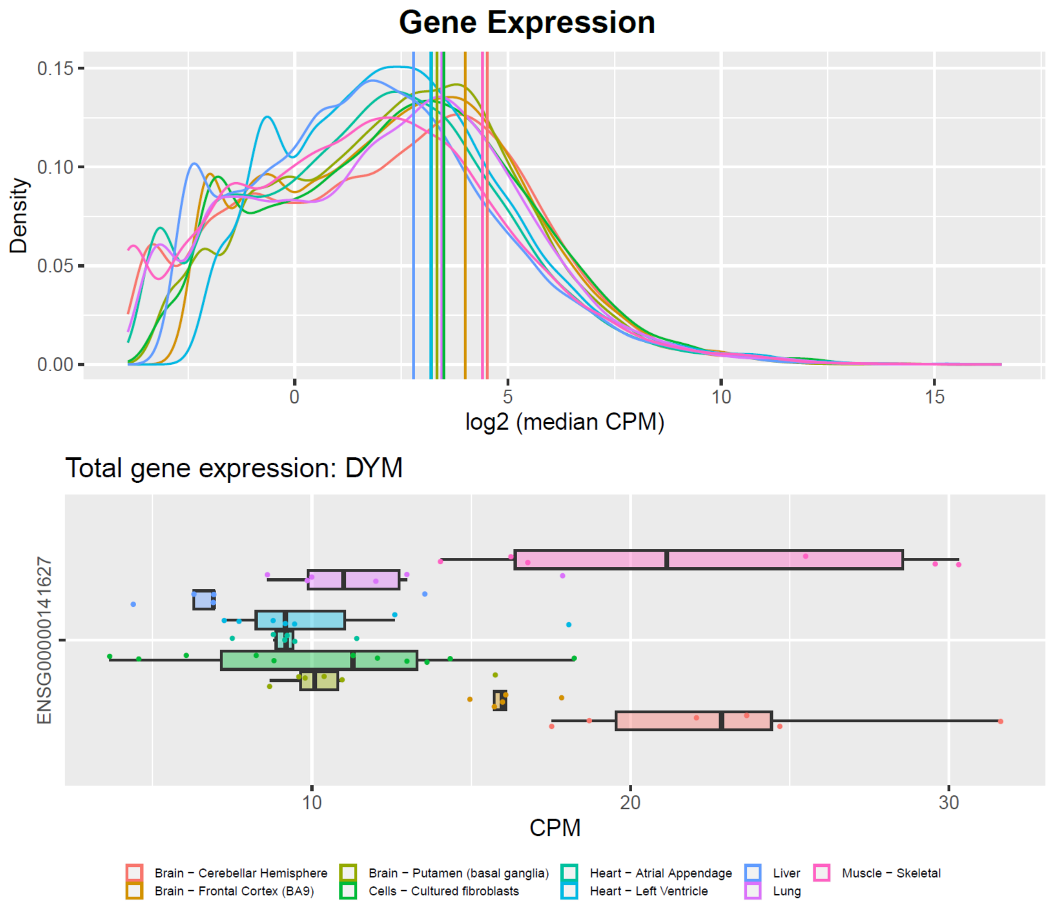


*Supplementary Figure 20: Gene expression of DYM.* Total gene expression of DYM across tissues. Density plot showing where the gene falls on the distribution of all genes expressed in each tissue.

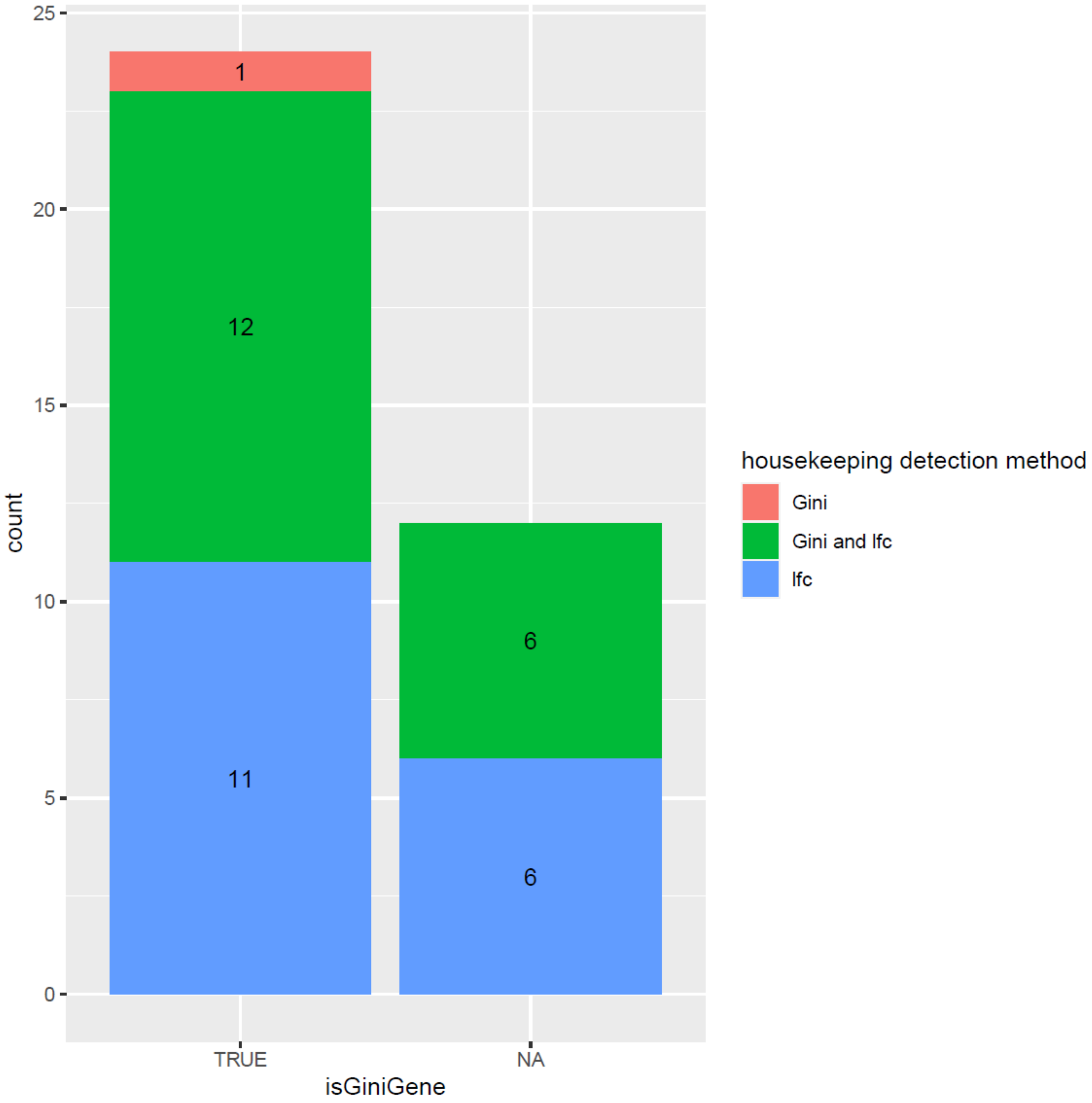


*Supplementary Figure 21: Showcasing the detection method we used for determining potential housekeeping isoforms.* We used the gini coefficient method (we chose Gini coefficient < 0.3 as our cutoff) as well as our original logfold change method. The X-axis shows whether the isoform we detected came from a gene that the Joshi et. al study identified as a Gini gene.

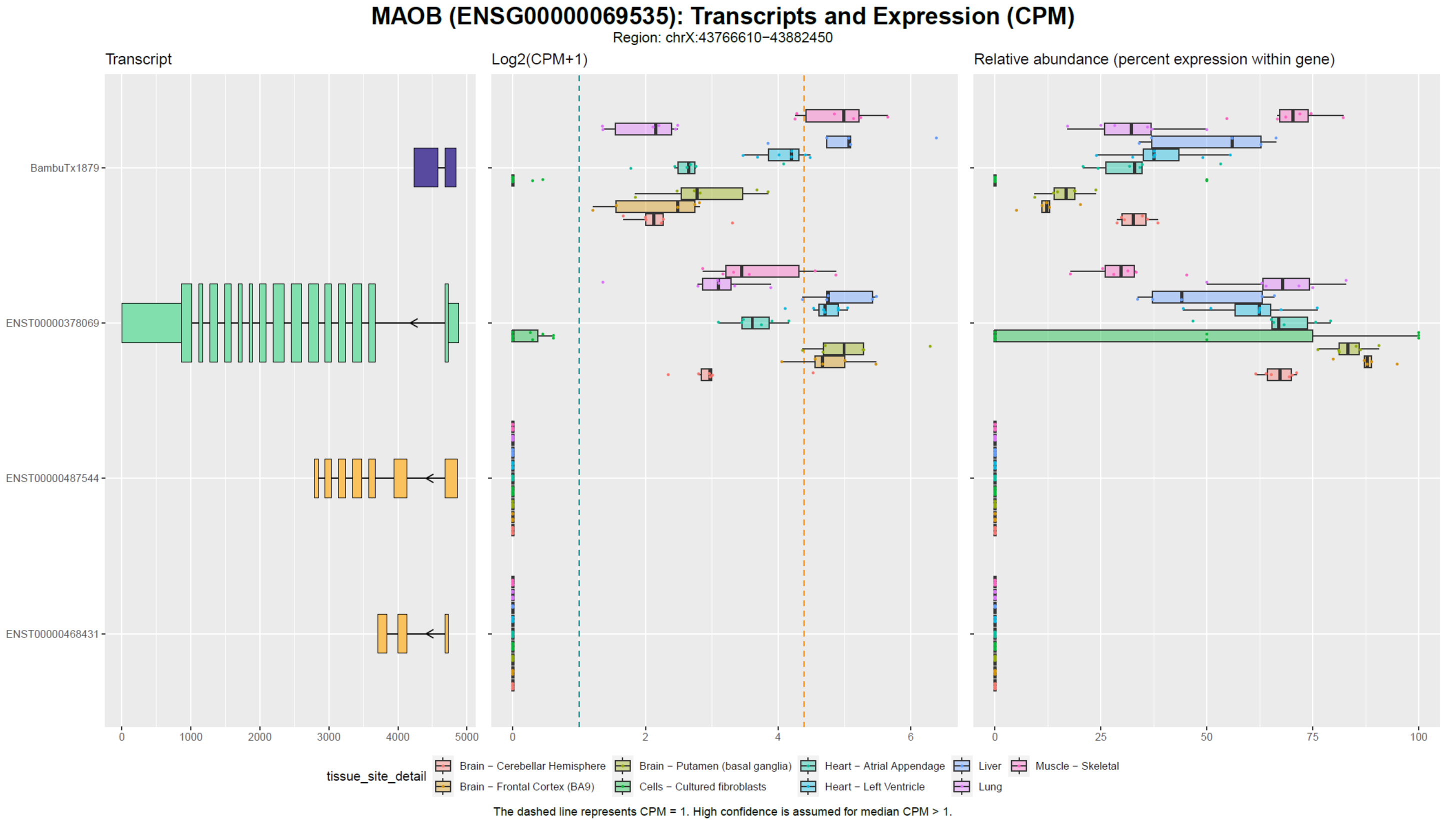


*Supplementary Figure 22: MAOB isoforms, expression, and relative expression*

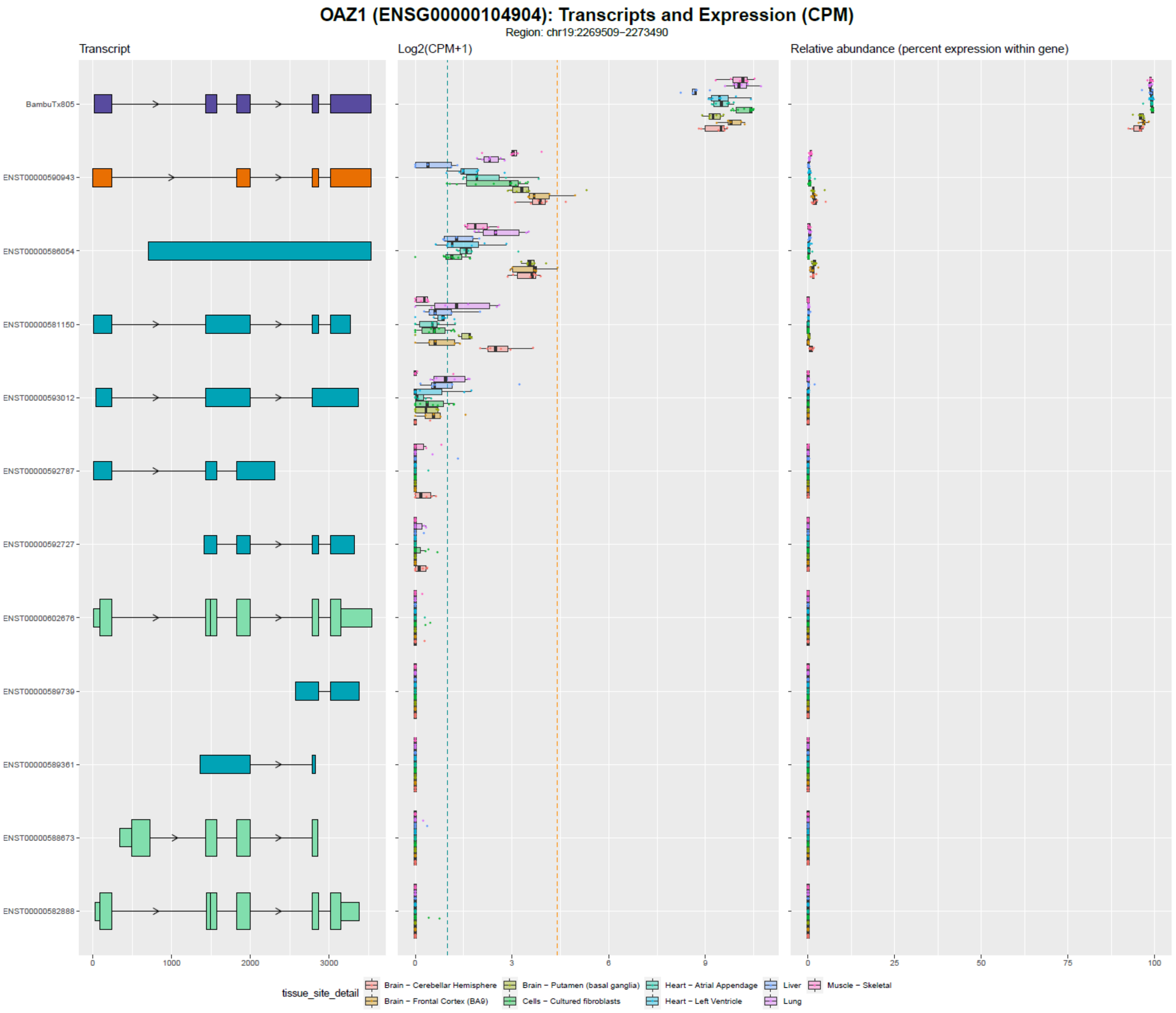


*Supplementary Figure 23: OAZ1 isoforms, expression, and relative expression*

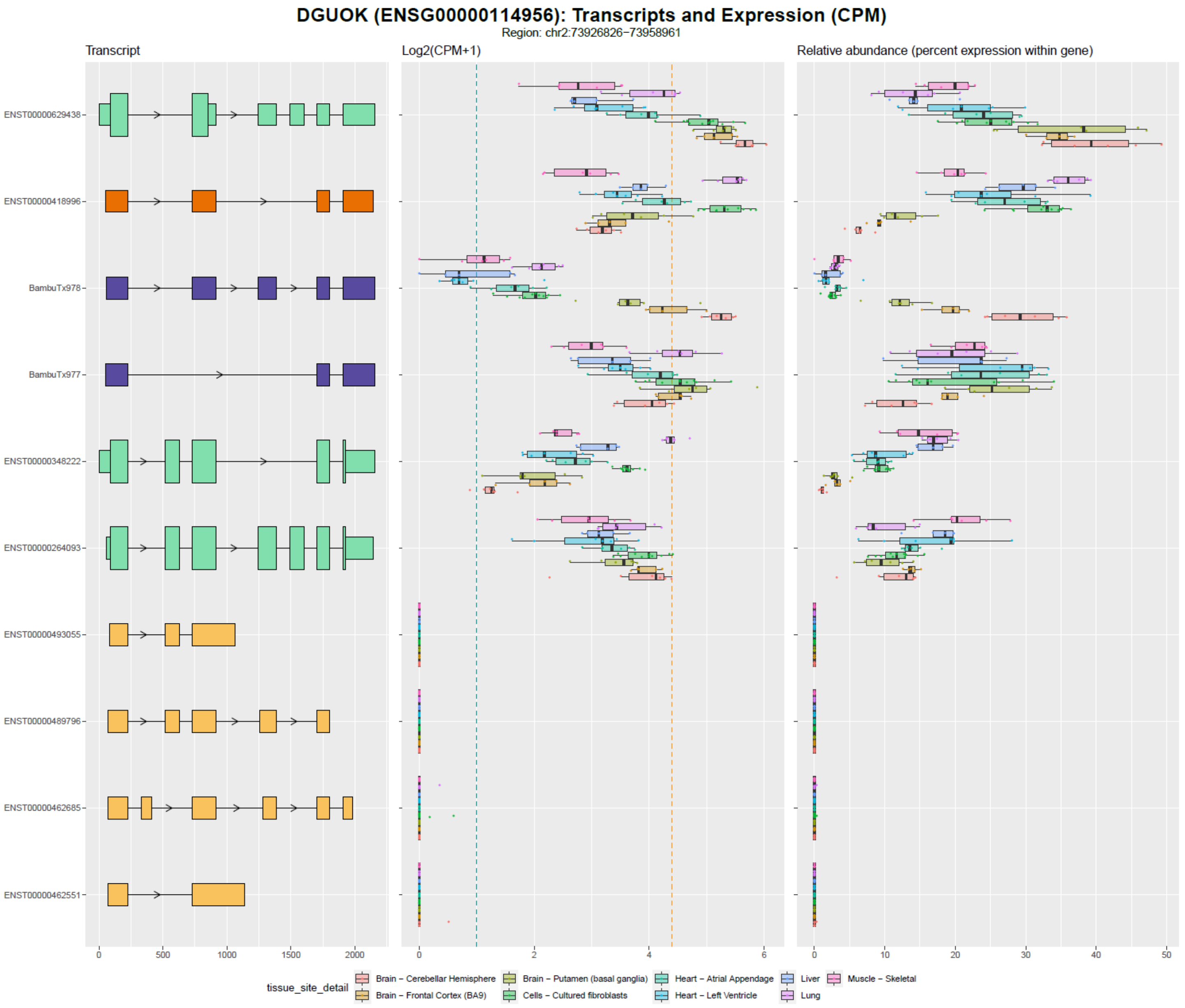


*Supplementary Figure 24: DGUOK isoforms, expression, and relative abundance*
